# Large-scale death of retinal astrocytes during normal development mediated by microglia

**DOI:** 10.1101/593731

**Authors:** Vanessa M. Puñal, Caitlin E. Paisley, Federica S. Brecha, Monica A. Lee, Robin M. Perelli, Emily G. O’Koren, Caroline R. Ackley, Daniel R. Saban, Benjamin E. Reese, Jeremy N. Kay

## Abstract

Naturally-occurring cell death is a fundamental developmental mechanism for regulating cell numbers and sculpting developing organs. This is particularly true in the central nervous system, where large numbers of neurons and oligodendrocytes are eliminated via apoptosis during normal development. Given the profound impact of death upon these two major cell populations, it is surprising that developmental death of another major cell type – the astrocyte – has rarely been studied. It is presently unclear whether astrocytes are subject to significant amounts of developmental death, or how it occurs. Here we address these questions using mouse retinal astrocytes as our model system. We show that the total number of retinal astrocytes declines by over 3-fold during a death period spanning postnatal days 5-14. Surprisingly, these astrocytes do not die by apoptosis, the canonical mechanism underlying the vast majority of developmental cell death. Instead, we find that microglia kill and engulf astrocytes to mediate their developmental removal. Genetic ablation of microglia inhibits astrocyte death, leading to a larger astrocyte population size at the end of the death period. However, astrocyte death is not completely blocked in the absence of microglia, apparently due to the ability of astrocytes to engulf each other. Nevertheless, mice lacking microglia showed significant anatomical changes to the retinal astrocyte network, with functional consequences for the astrocyte-associated vasculature leading to retinal hemorrhage. These results establish a novel modality for naturally-occurring cell death, and demonstrate its importance for formation and integrity of the retinal gliovascular network.

## Introduction

Naturally-occurring developmental cell death contributes to the histogenesis of most tissues [1,2]. For example, in the mammalian central nervous system (CNS), many populations of neurons are subject to large-scale death that eliminates as many as half of the cells originally produced during neurogenesis [3,4]. Cell removal on this scale has a profound impact on neuroanatomy and circuit structure, and is essential for key steps in CNS morphogenesis [5–8]. Thus, to understand the fundamental developmental mechanisms that sculpt circuit anatomy and function, it is critical to document the extent of naturally-occurring CNS death, and the mechanisms by which it occurs.

In mammals, naturally-occurring cell death typically occurs via apoptosis, during specific developmental periods [4]. Following death, professional phagocytes such as CNS microglia are recruited to clear apoptotic corpses. This sequence of events is critical for normal development, since mice lacking essential apoptotic genes exhibit a range of defects affecting the architecture of the CNS and the patterning of its cellular populations [4,5,9,10]. Autophagy and necroptosis are non-apoptotic mechanisms that can occur in a variety of contexts, but their contribution to mammalian CNS development is minimal [11,12]. However, death by other mechanisms could still make important contributions to CNS development [13].

It has recently emerged that CNS cell death can be mediated through engulfment and killing of viable cells [14]. In contrast to apoptosis, where phagocytosis serves a debris-removal role, this variety of death involves phagocytosis prior to or concomitant with cell killing. As such, it may be termed “death by phagocyte.” Microglia have been shown to engage in phagocytic killing in pathological contexts [15–17] and in the developing mammalian forebrain, where they appear to eliminate live neural progenitors within the subventricular zone [18]. These observations raise the possibility that death by phagocyte might sculpt other developing CNS cell populations. However, this possibility remains to be tested.

While developmental death of neurons has been extensively studied for almost a century, few studies have examined the role of death in the development of astrocytes. Because astrocytes have many essential roles in neuronal and vascular physiology [19–22], it is critical to understand the histogenetic mechanisms – such as regulation of cell number – that shape astrocyte development. Apoptotic astrocytes have been reported in developing cortex, retina, and cerebellum [23–29], but there is little evidence that such death is extensive enough to change astrocyte number. It remains to be determined whether death sculpts the developing astrocyte population, and if so, what the death mechanism might be.

To investigate the scale of death during astrocyte development, we chose as our primary model system the nerve fiber layer astrocytes of mouse retina. This astrocyte population resides at the retinal surface, in close association with blood vessels and retinal ganglion cell (RGC) axons (Fig 1A). We chose these cells for two reasons. First, they are confined to a monolayer within the retinal nerve fiber layer (RNFL), simplifying estimates of absolute cell number – a crucial advantage for studies of cell death. Second, regulation of astrocyte number may have implications for retinal function and disease. During angiogenesis, endothelial cells colonize the retina by using astrocyte arbors as an angiogenic patterning template [22]. When astrocyte numbers are experimentally elevated or lowered, both astrocyte and vascular patterning become disturbed, impairing vascular integrity [30–32]. It is therefore likely that astrocyte numbers are tightly developmentally regulated to prevent vascular pathology.

**Figure 1.**
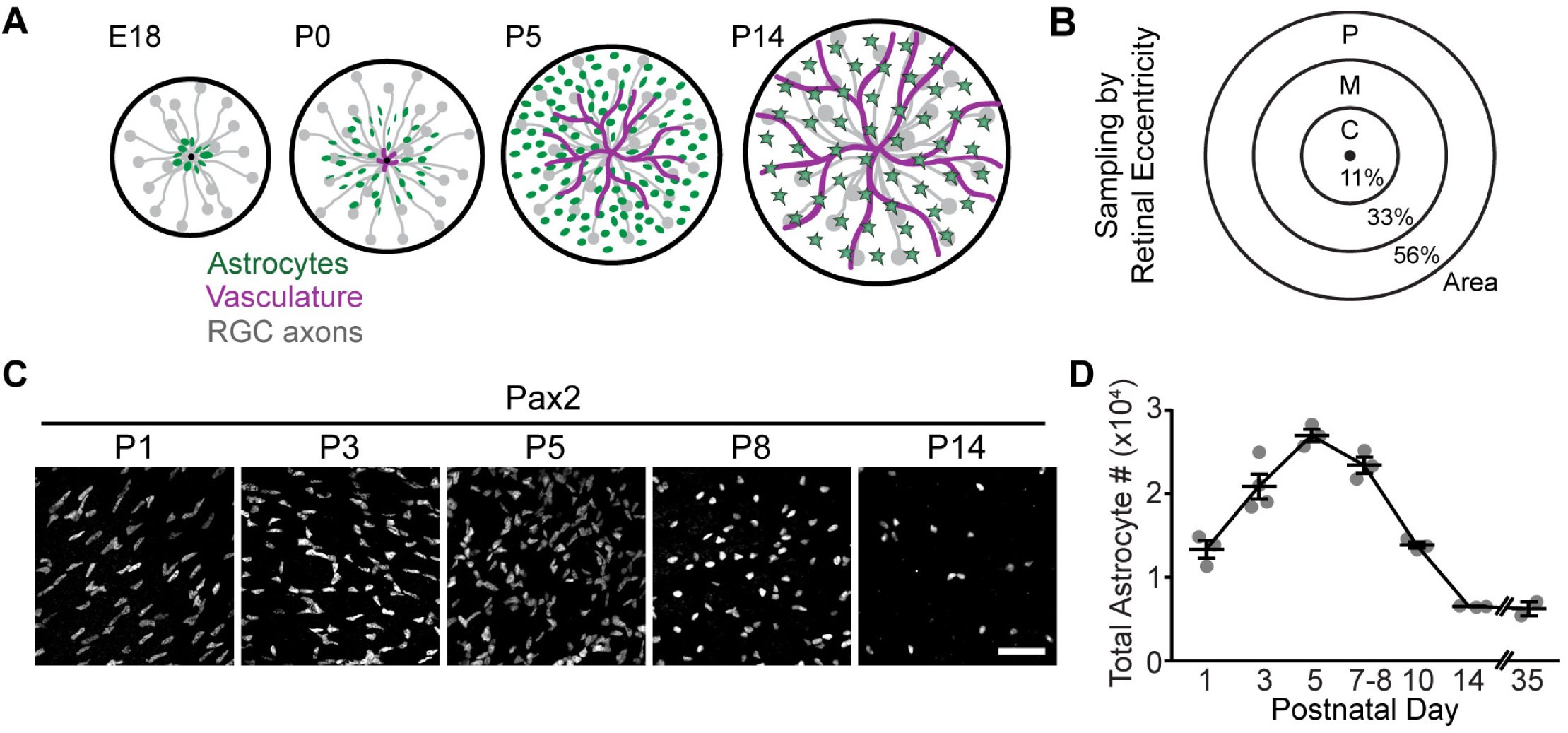
Developmental changes in astrocyte population size due to cell death. **A**) Schematic of mouse RNFL development showing timing of retinal colonization by astrocytes and vasculature. Stars denote mature astrocytes. **B**) Method to quantify total astrocyte number. Weighting was according to fraction of retinal area within each sampled region (C, central; M, middle; P, peripheral). **C**) Astrocyte density, shown in en-face images from retinal whole-mounts stained for Pax2. **D**) Quantification of total astrocyte numbers across development, using retinas stained for Pax2 or Sox9 (see S1 Fig. A,B). Error bars: mean ± S.E.M. Sample sizes depicted by data points on graph. Scale bar = 50 µm.

We find that retinal astrocytes are initially overproduced and then culled during a brief period of postnatal development. Surprisingly, apoptosis is not a major driver of their death; instead, we demonstrate that death by phagocyte plays a key role in this process. During the astrocyte death period, microglia interact extensively with astrocytes and engulf astrocytic material. These microglial behaviors are related to astrocyte killing, rather than mere clearance of already-dead corpses, because ablation of microglia *in vivo* increased the number of viable, fully differentiated astrocytes. Finally, we find that astrocytic death by phagocyte can occur through either a heterotypic or a homotypic mechanism: When microglia are absent, astrocytes can partially compensate for their loss by eliminating each other. Together these data reveal that retinal astrocyte death has important mechanistic differences from other developmental deaths in the CNS. Thus, death by phagocyte might have broader roles in tissue morphogenesis than previously appreciated.

## Results

### Developmental changes in astrocyte population size due to cell death

To investigate whether retinal astrocytes have a period of naturally-occurring cell death, we estimated total astrocyte number across postnatal mouse development. Estimates were generated from retinas stained in whole-mount for nuclear astrocyte markers Pax2 or Sox9 (Fig. 1C; S1 Fig. A,B; [28]) using a sampling strategy described previously ([30]; Fig. 1B). This analysis showed that retinal astrocyte numbers increased until P5, then declined substantially – by over 3-fold – to reach adult levels by P14 (Fig.1C,D). The rising phase of this curve is predominantly due to astrocyte migration into the retina, which is complete by P4-5 ([22]; Fig. 1A). Astrocyte loss was not due to migration out of the tissue; nor was it due to transdifferentiation into a non-astrocyte cell type, as shown by Cre-lox lineage tracing (S1 Fig. C,D). Moreover, the cell loss cannot be ascribed simply to downregulation of the early astrocyte marker Pax2 [28], as Sox9 gave identical results (S1 Fig. A,B). These results rule out the most likely alternative explanations for astrocyte disappearance, leading us to conclude that cell death is responsible.

### Retinal astrocyte death is not due to apoptosis

Since apoptosis is typically the mechanism for developmental cell death, we tested whether this was the case for retinal astrocytes. To this end we stained retinas with antibodies to cleaved-caspase 3 (CC3), a histological marker of apoptotic cells [33]. Unexpectedly, few developing astrocytes expressed CC3 (Fig. 2A; S2 Fig. C; *n*=14 CC3^+^ astrocytes out of 29,383 analyzed across all ages). Therefore, astrocyte apoptosis was quite rare.

**Figure 2.**
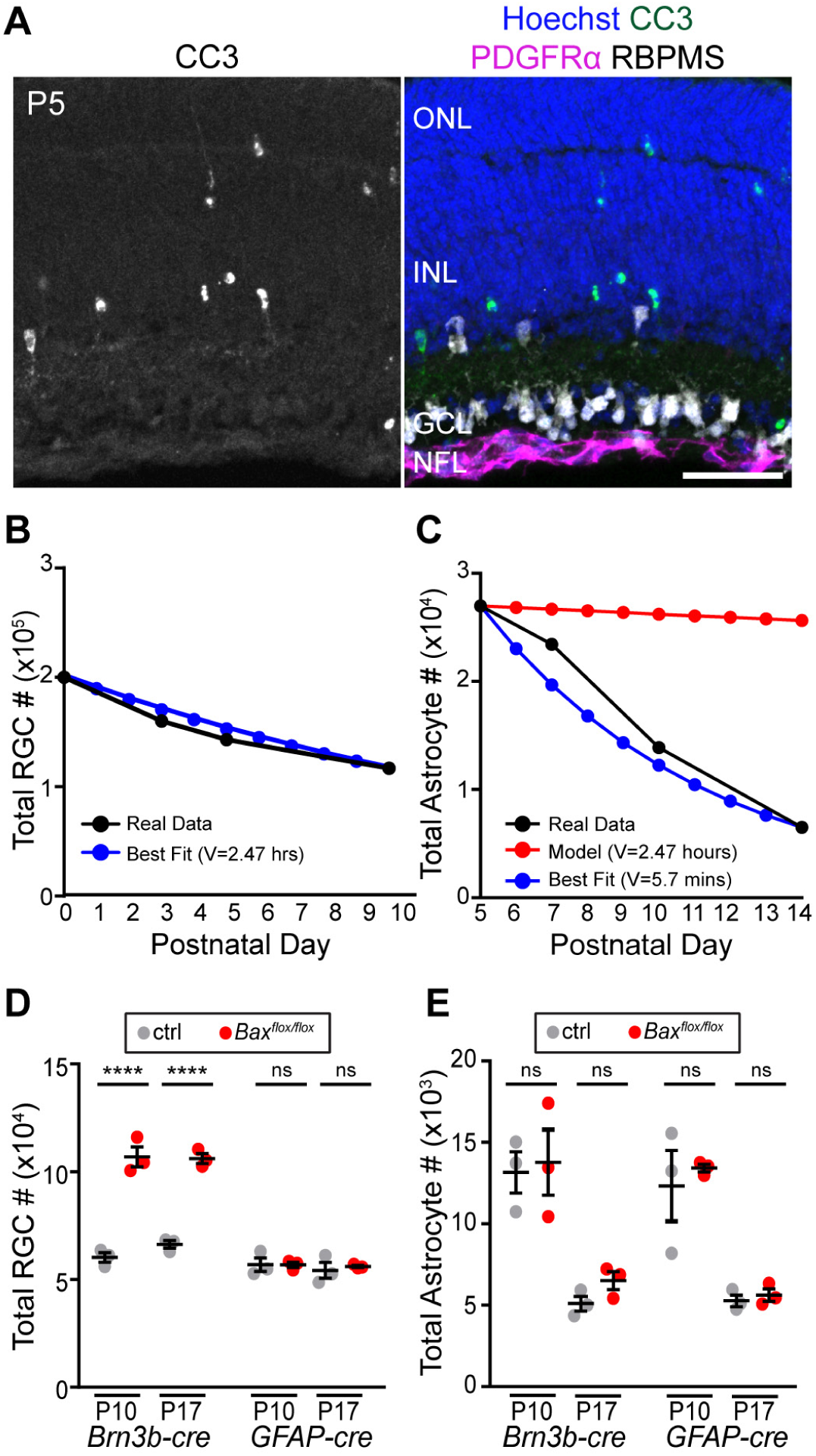
Retinal astrocyte loss is not due to apoptosis. **A**) P5 retinal sections stained for CC3 (apoptosis marker), PDGFRα (astrocyte cell surface marker), RBPMS (RGC marker), and Hoechst (nuclear stain). CC3^+^ cells are frequently found in neuronal layers, but not in retinal nerve fiber layer (RNFL) where astrocytes reside. **B,C**) Modeling to determine effect of observed apoptosis rate on RGC (B) and astrocyte (C) population size. Black: Observed cell number (B, RGC data from [36], see also S2 Fig. D; C, astrocyte numbers from Fig. 1D). Blue: Best fit models using observed apoptotic cell fraction as death rate parameter (B, [36]; C, CC3^+^ astrocyte fraction from S2 Fig. C). Clearance time parameter in best fit model is given. Red in C: Astrocyte model with clearance time parameter set to biologically plausible value (2.47 h); apoptosis alone cannot account for observed astrocyte numbers. **D,E**) Effects of cell-type-specific *Bax* deletion on RGC (D) and astrocyte (E) numbers. RGCs, *Brn3b-Cre*; astrocytes, *GFAP-Cre*. Statistics: For each cell type, 3-way ANOVA followed by Tukey’s post-hoc test; ****p<0.0001. NS P-values ≥ 0.99. N=3 for all groups. Error bars: mean ± S.E.M. Sample sizes depicted by data points on graph. Scale bar: 50 µm.

Apoptotic corpses can be cleared in a matter of hours, such that only a small fraction of the dying population is visible in any given histological sample [34,35]. To ask whether the small number of CC3^+^ astrocytes could reflect such a large change in cell number, we took a modeling approach. The model we used [35] was developed to estimate apoptotic cell clearance time, given experimental measurements of cell numbers and the fraction of visibly dying (i.e. CC3^+^) cells. If the clearance time is known, the model can also be used to calculate the expected cell number decline given a measurement of the CC3^+^ dying cell fraction. To validate this approach, we first modeled the developmental decline in RGC numbers previously reported in rat retina [36] (see Methods). The model’s best-fit curve predicted RGC clearance time as 2.47 hours (Fig. 2B), which is in alignment with published clearance times of ∼1-3 hours for apoptotic neurons and oligodendrocytes [37–40]. We therefore applied the model to our astrocyte data. According to the model, in order for the observed percentage of apoptotic astrocytes to account for the observed cell loss, astrocyte clearance would need to occur in an implausibly short time – under 6 min (Fig. 2C). When the RGC clearance time was used (2.47 hours), the model predicted only a small decline in astrocyte number by P14, nowhere near the measured value (Fig. 2C). These results strongly suggest that the observed frequency of astrocyte apoptosis is too low to plausibly account for the P5-P14 decline in cell number.

Given the scarcity of astrocyte apoptosis, we hypothesized that astrocyte death would proceed normally when apoptosis is genetically perturbed. *Bax* is an essential apoptotic gene, particularly in the CNS, where loss of its function impairs neuronal death and increases neuron numbers [14]. Using a conditional *Bax*^*flox*^ allele, we confirmed previous reports [41,42] that deletion of *Bax* in RGCs rescues them from death (Fig. 2D). By contrast, deletion of *Bax* using a *GFAP-cre* line, which is highly selective for astrocytes and Müller glia [30] (S1 Fig. D), did not prevent astrocyte death (Fig. 2E). Thus, astrocytes die in a *Bax*-independent manner. Taken together, our CC3 and *Bax* studies demonstrate that apoptosis is not primarily responsible for astrocyte developmental death.

### Microglia engulf retinal astrocytes during their death period

We next investigated non-apoptotic death programs that might eliminate astrocytes. Electron micrographs of the developing RNFL showed no histological hallmarks of autophagy or necroptosis [33], arguing against involvement of these mechanisms (S2 Fig. B). To explore a role for microglia, we began by examining their localization and morphology, using *Cx3cr1*^*CreER-ires-YFP*^ mice [43] (abbreviated *Cx3cr1*^*CreER*^). Microglia preferentially accumulated in the RNFL during the astrocyte death period (Fig. 3A,B), and sent out processes contacting the vast majority of astrocytes (Fig. 4D). Furthermore, during the death period, RNFL microglia exhibited both morphological and molecular signatures of phagocytic activity. Morphologically, they had an amoeboid shape typical of phagocytic macrophages [44] (Fig. 3C). Molecularly, they selectively expressed CD68 and Osteopontin, two markers of phagocytic microglia [45,46] (Fig. 3D-F). Once the death period was over, however, microglia assumed a resting morphology and downregulated both markers (Fig. 3C-G). Together, these observations demonstrate that phagocytic microglia are present at the right time and place to play a role in developmental astrocyte death.

**Figure 3.**
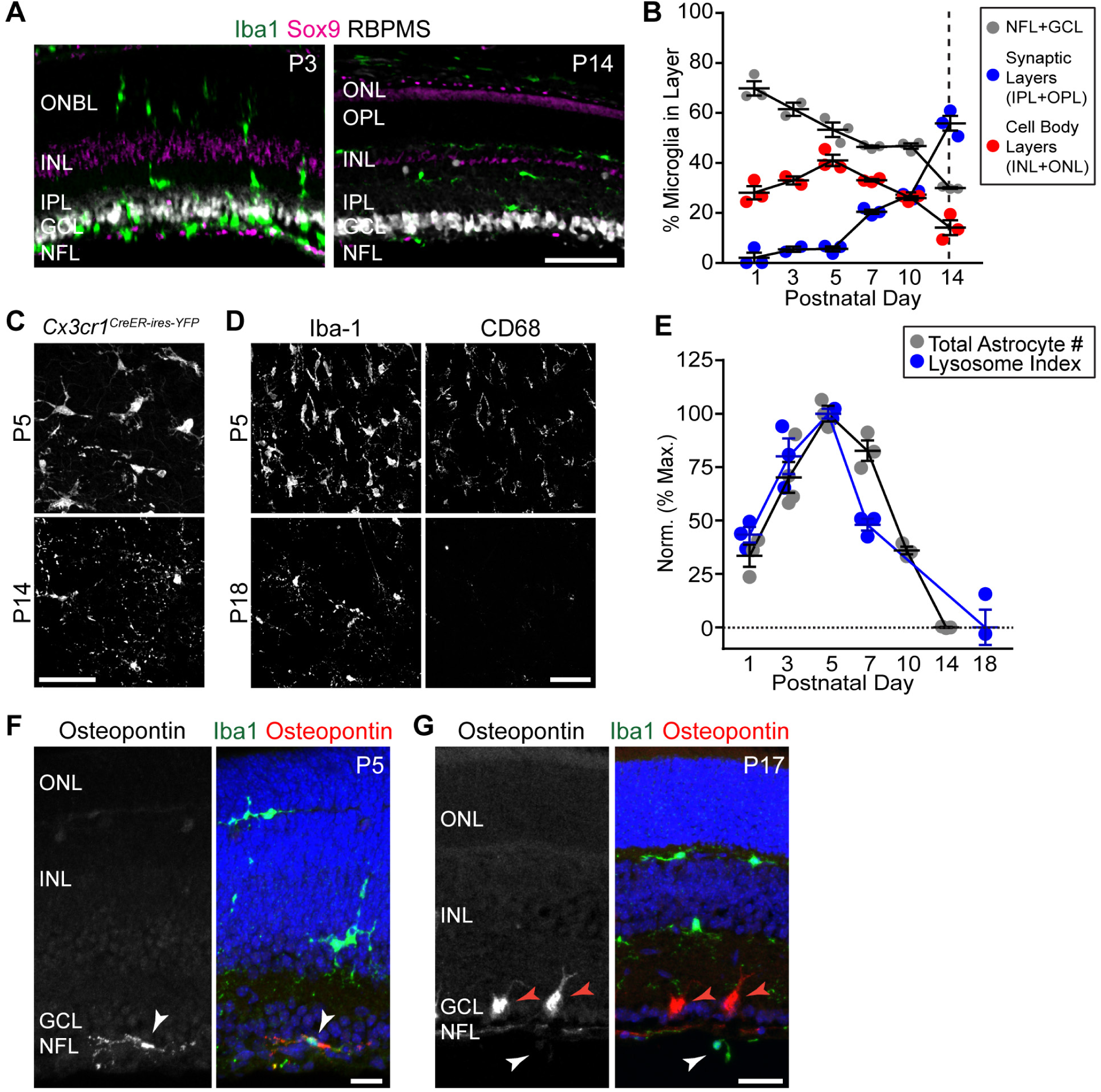
Nerve fiber layer microglia express a phagocytic phenotype during the astrocyte death period. **A,B**) Retinal microglia localize preferentially to RNFL until end of astrocyte death period at P14 (dashed line). A: Representative retinal sections showing laminar location of Iba1^+^ microglia; B: Microglial laminar location quantified across development in whole-mount images (e.g. C). **C**) RNFL microglia visualized in *Cx3cr1*^*CreER-ires-YFP*^ whole-mount using anti-GFP. Microglia show amoeboid morphology during astrocyte death period (P5) but ramified morphology once death is complete (P14). **D**) RNFL microglia (labeled with anti-Iba1) have high lysosomal content (anti-CD68) at peak of astrocyte number (P5). CD68 is largely absent after the death period (P18). **E**) Quantification of microglial phagocytic capacity, measured using a lysosome index (CD68^+^ lysosome content per microglial cell; blue). Gray: Total astrocyte numbers, replotted from Fig. 1D for comparison. Note strong correlation between these measures, both during arrival of astrocytes in retina (P1-5; see Fig. 1A) and during their elimination (P5-14). **F,G**) During death period, RNFL microglia (white arrows) are molecularly distinct from Iba1^+^ microglia in other layers, as shown by their expression of Osteopontin at P5 (F) but not P17 (G). Red arrows: Osteopontin^+^ RGCs [67]. Abbreviations: NFL, Nerve Fiber Layer; GCL, Ganglion Cell Layer; IPL, Inner Plexiform Layer; INL, Inner Nuclear Layer; ONBL, Outer Neuroblast Layer; OPL, Outer Plexiform Layer; ONL, Outer Nuclear Layer. Error bars: Mean ± S.E.M. Sample sizes depicted by data points on graph. Scale bars: 50 µm (A,C,D); 20 µm (F,G).

**Figure 4.**
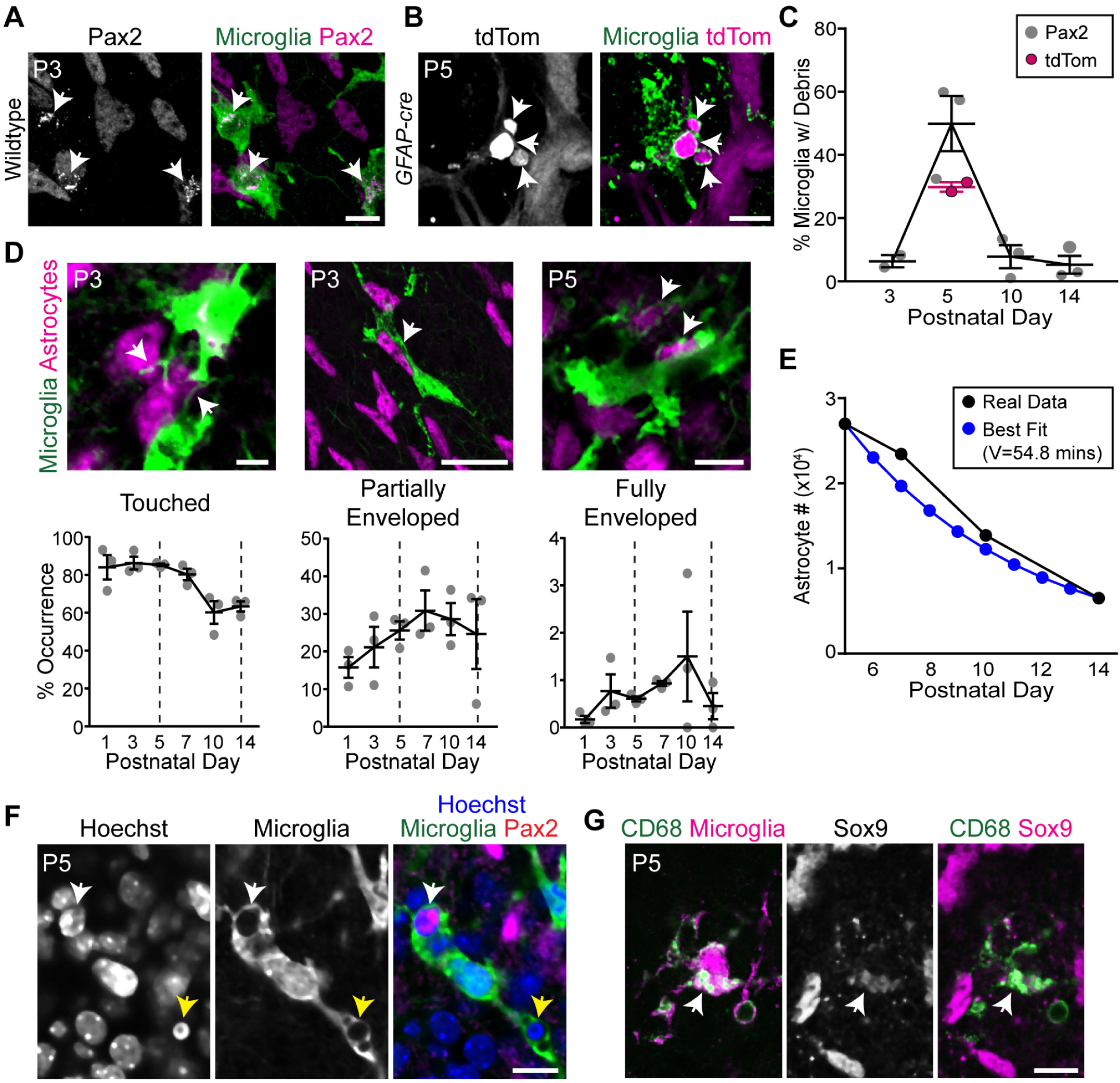
Microglia engulf developing retinal astrocytes. **A,B**) Representative images showing astrocyte debris within microglia (arrows). Astrocytes labeled with antibody against Pax2 (A) or with tdTomato (tdTom) Cre reporter driven by *GFAP-cre*(B). This Cre line is selective for astrocytes at P5 and is not yet expressed by Müller glia [30]. Microglia labeled with anti-GFP in *Cx3cr1*^*CreER-ires-YFP*^mice (A) or antibodies to Iba1 and P2Y12 (B). A: single optical plane; B: Z-projection of 3 optical planes (1.2 µm Z distance total). **C**) Percentage of microglia containing astrocyte debris peaks at P5, coinciding with peak astrocyte number. **D**) Astrocyte-microglia interactions quantified in *Cx3cr1*^*CreER-ires-YFP*^mice. Bottom row: Percentage astrocytes touched, partially enveloped, or fully enveloped by microglial processes. Examples of each interaction category are shown (top row; arrows). Dashed lines: start (P5) and end (P14) of astrocyte loss. Phagocytosis-like interactions (i.e. partially or fully enveloped astrocyte somata) peak during death period. **E**) Modeling suggests engulfment rate is sufficient to account for developmental changes in astrocyte population size. Black: Observed astrocyte number from Fig. 1D. Blue: Best fit model using average frequency of engulfment determined in (D) (“fully enclosed” category) as death rate parameter. Clearance time parameter of best fit model (54.8 min) is plausible given past studies [18,47]. **F**) Engulfed astrocytes do not have pyknotic nuclei. White arrow: non-pyknotic astrocyte nucleus within a microglial phagocytic cup. Yellow arrow: non-astrocytic pyknotic nuclei within phagocytic cup of the same microglial cell. Single optical plane shown. Microglia labeled with *Cx3cr1*^*CreER-ires-YFP*^. **G**) Astrocyte debris (arrow) within CD68^+^ lysosomal compartments of *Cx3cr1*^*CreER-ires-YFP*^ microglia. All images are *en-face*views of whole-mounted retina. Error bars, mean ± S.E.M. Sample sizes depicted by data points on graphs. Scale bars: 5 µm (D, left); 10 µm (D, right; F; G); 25 µm (A; B; D, center).

To test whether the phagocytic state of RNFL microglia reflected engulfment of astrocytes, we analyzed retinal tissue double-labeled for each cell type. Astrocyte nuclear debris, immunopositive for Pax2 or Sox9, was found within microglial intracellular compartments (Fig.4A,C,G; S1 Movie). These compartments co-labeled for CD68, consistent with the expected lysosomal destination of engulfed material (Fig. 4G). Internalization of astrocyte debris was also confirmed using tdTomato as a genetically encoded astrocyte label (Fig. 4B,C). Further, we observed microglial processes that partially or fully enveloped astrocyte somata, suggesting they were being engulfed (Fig. 4D; S2 Movie; S3 Movie). These processes often resembled the classic phagocytic cups that surrounded pyknotic neuronal corpses (Fig. 4F). Notably, the nuclei of engulfed astrocytes were not pyknotic or degenerated (Fig. 4F), suggesting they may be engulfed while still viable. Anatomical signatures of astrocyte engulfment were detected most frequently during the period of astrocyte death, suggesting that engulfment occurred selectively during this time (Fig. 4C,D). Therefore, the phagocytic state of RNFL microglia during the death period is due at least in part to their engulfment of astrocytes.

To ask whether engulfment occurs often enough to explain the decline in astrocyte numbers, we once again took a modeling approach. Using the same model as in our apoptosis studies (Fig. 2C), we calculated astrocyte clearance time based on the observed average number of “fully enveloped” astrocytes (Fig. 4D). The model (Fig. 4E) predicted a clearance time of 54 min, closely matching previous reports of the time needed for microglia to clear engulfed cells (∼45 min) [18,47]. Therefore, astrocyte engulfment occurs with sufficient frequency to plausibly account for astrocyte death.

### Blockade of major phagocytosis pathways does not impact astrocyte number

The results so far led us to hypothesize that microglial phagocytosis is responsible for causing astrocyte death. To test this possibility, we first sought to prevent microglia from engaging in phagocytosis. Microglia express several well-characterized receptors that are required for engulfment of dead cells, cellular debris, and in some cases even viable cells [16,48–52]. We reasoned that one of these pathways might also mediate elimination of developing astrocytes. We therefore examined mutant mice in which the best characterized receptors or their downstream signaling components were eliminated. These included mutants lacking the complement receptor CR3 (*Itgam*^*–/–*^); the Mer tyrosine kinase receptor (*Mertk*^*–/–*^); and Syk tyrosine kinase (*Cx3cr1*^*CreER*^; *Syk*^*flox/flox*^), an essential signal downstream of FC-gamma and TREM-2 [51,53]. If these pathways are required for astrocyte elimination, astrocyte numbers should be increased in mutants. However, none of the mutants showed such an effect (Fig. 5A-C).

**Figure 5.**
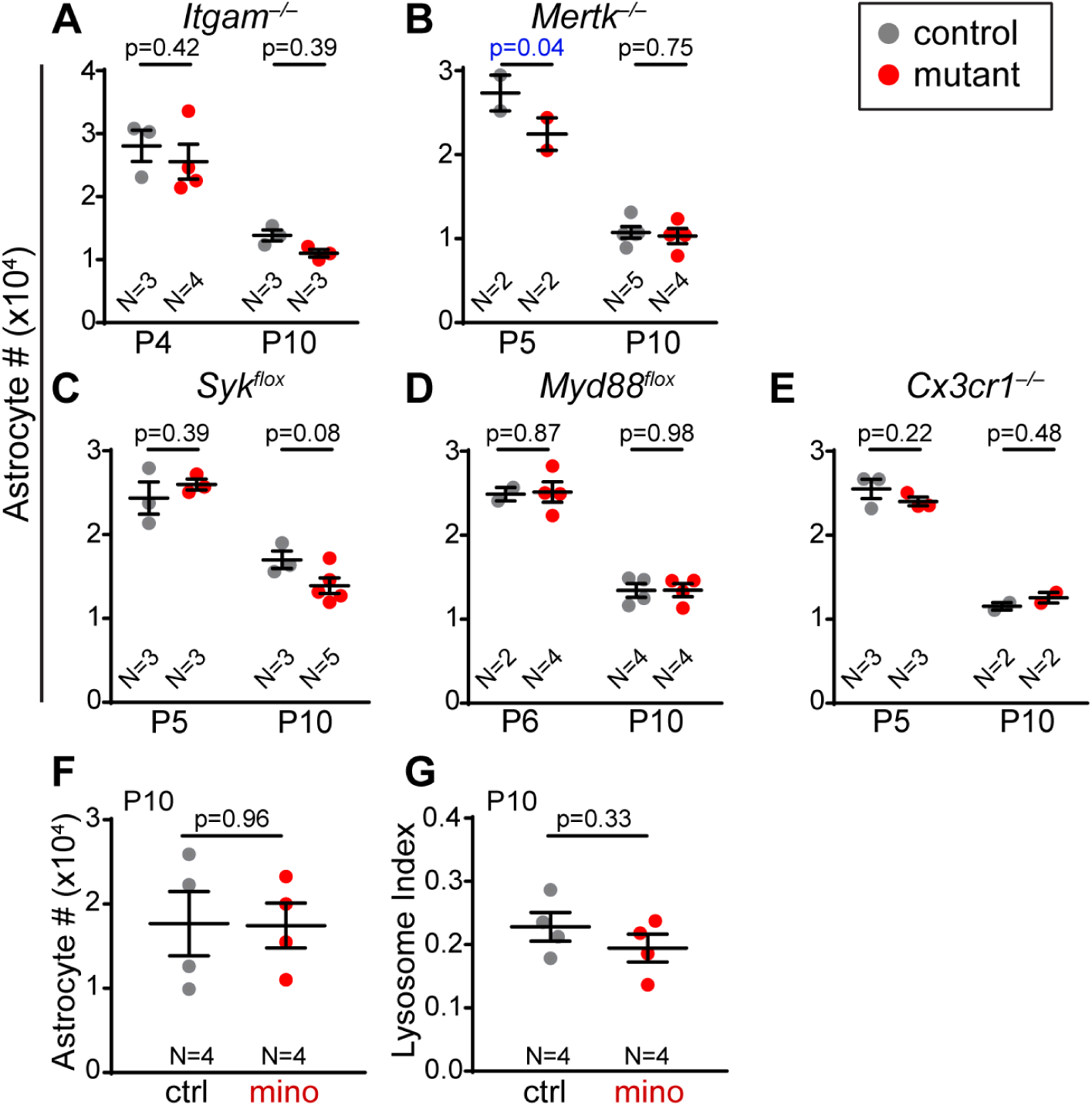
Blockade of major phagocytosis pathways does not affect astrocyte number. **A-E**) Total astrocyte numbers were determined in mutant mice (red) and littermate controls (gray) at two ages during astrocyte death period. A: *Itgam* constitutive null mutants lack complement C3 receptor. B: *Mertk* constitutive null mutants lack Mer tyrosine kinase receptor. C,D: *Syk*^*flox*^(C) and *Myd88*^*flox*^(D) are conditional alleles of signaling components downstream of FcγR/TREM2 (C) or toll-like receptors (D). Mutants carried *Cx3cr1*^*CreER*^ (C) or *Cx3cr1-Cre* (D) to drive microglia-specific gene deletion. E: *Cx3cr1*^*CreER/CreER*^ constitutive null mice lack fractalkine receptor. Statistics: For each data set, 2-way ANOVA followed by Fisher’s LSD post-hoc test. Blue, p-value < α. **F,G**) No effect of minocycline on total astrocyte numbers (F) or microglial phagocytic capacity (G). Administration, P4-P9; analysis, P10. Statistics: Two-tailed t-tests. Error bars: Mean S.E.M.

We next investigated the role of two well-characterized signaling pathways that are potent regulators of microglial physiology, including their phagocytic activity. First, we tested the role of the pro-inflammatory Toll-like receptors using a microglia-specific deletion of their essential downstream signaling molecule MyD88 (*Cx3cr1-Cre; Myd88*^*flox/flox*^). Second, we examined mice lacking the CX3CR1 receptor (*Cx3cr1*^*CreER/CreER*^). Neither mutant showed an astrocyte survival phenotype (Fig. 5D,E). Finally, we tested minocycline, a drug that suppresses inflammation and microglial phagocytic activity through an unknown mechanism. This too failed to affect microglial phagocytic activity, as measured by both CD68^+^ lysosome content and astrocyte numbers (Fig. 5F,G). Taken together, these experiments indicate that the best-characterized pro-inflammatory and pro-engulfment microglial pathways are either not involved in astrocyte death, or they are capable of completely compensating for each other when one is absent.

### Ablation of microglia increases astrocyte number

One possible interpretation of the receptor mutant studies is that microglia mediate astrocyte removal through a new pathway, or through a combination of the tested pathways. However, it also remains possible that microglia are not in fact involved in astrocyte developmental death. To distinguish between these possibilities, we ablated microglia during the death period – starting at P4, and continuing until the end of the death period at P14. If microglia are responsible for culling the astrocyte population, then ablating microglia during this period should prevent astrocyte death.

To ablate microglia, we used an established chemogenetic strategy [43] whereby *Cx3cr1*^*CreER*^ drives microglia-specific expression of a tamoxifen-inducible diphtheria toxin receptor (DTR). Because these tools had not previously been used during retinal development, we first confirmed their specificity, efficacy, and temporal characteristics (S3 Fig.). Based on these experiments we developed a protocol for repeated tamoxifen and diphtheria toxin administration that selectively eliminated microglia as early as P6 and prevented their return for the entirety of the death period (Fig. 6A,B).

**Figure 6.**
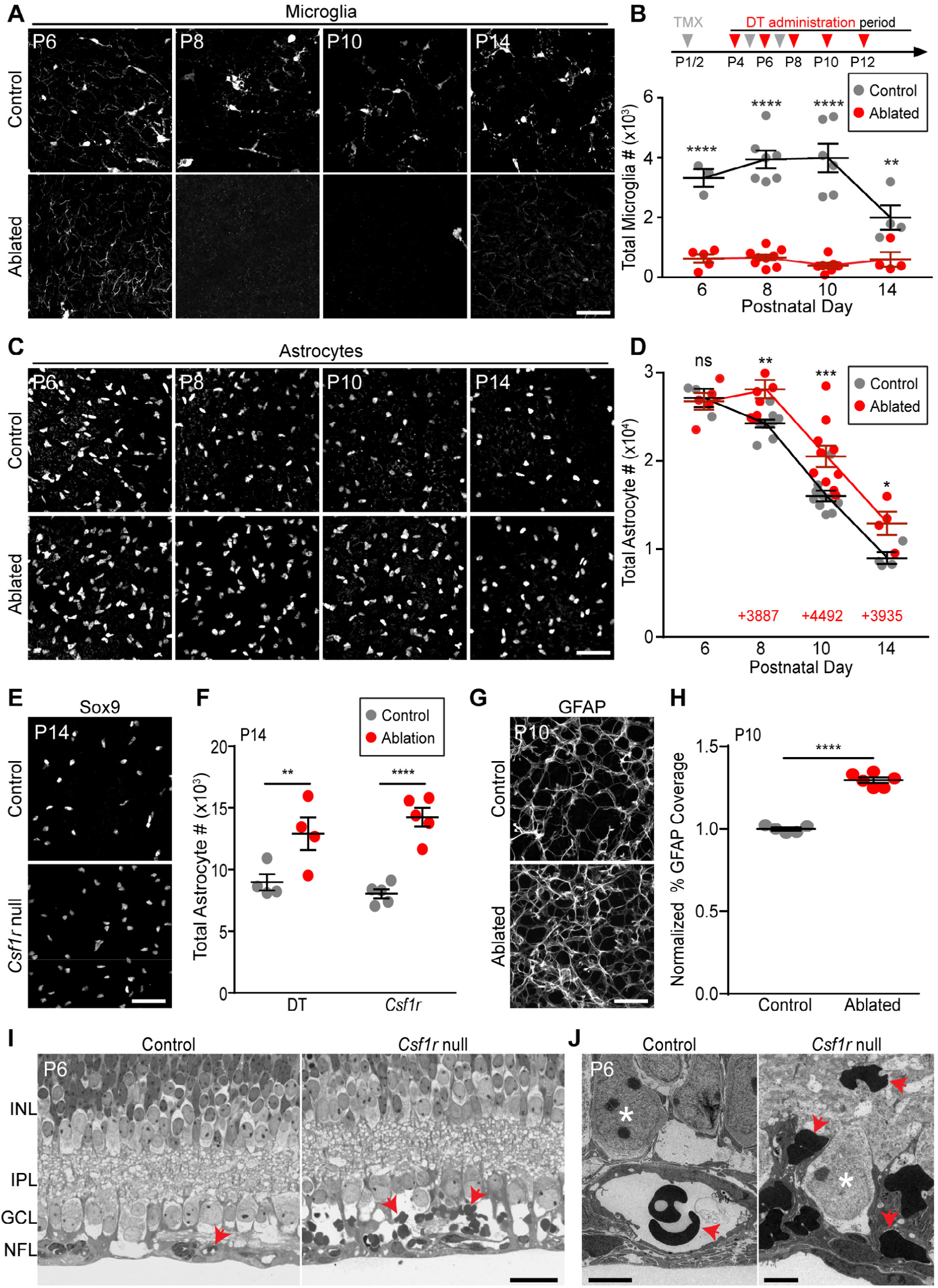
Ablation of microglia increases astrocyte number. **A,B**): Microglial ablation paradigm. Tamoxifen (TMX, gray arrows) and diphtheria toxin (DT, red arrows) were administered at indicated times (B, top). A: representative confocal images of whole-mount *Cx3cr1*^*CreER-ires-YFP*^retina showing microglial morphology (anti-GFP). B: Quantification of microglial numbers. Ablated mice carried both *Cx3cr1*^*CreER*^and *Rosa26*^*iDTR*^transgenes; littermate controls lacked one of the transgenes. Statistics: 2-way ANOVA followed by Fisher’s LSD post-hoc test. ****P<0.0001; **P=0.0037. **C,D**) Effects of microglial ablation on astrocyte number. C: Representative images of Pax2^+^ or Sox9^+^ astrocytes in control and microglia-ablated animals. D: Quantification of astrocyte number. Red text: number of excess astrocytes in ablated animals at each age. Statistics as in B. *P=0.0374; **P=0.0021; ***P=0.0003; ns=0.8439. **E,F**) Elimination of microglia using *Csf1r* null mice. Increase in total astrocyte number at P14 was similar to DT ablation paradigm. Statistics: 2-way ANOVA followed by Fisher’s LSD post-hoc test. **P=0.0054; ****P<0.0001. **G,H**) Increased density of GFAP^+^ astrocyte network in ablated retina. G: Representative en-face confocal images of RNFL. H: Quantification of area covered by GFAP+ arbors, from images similar to G. Statistics: Two-tailed t-test (P<0.0001). **I,J**) Thin plastic sections of *Csf1r* null mutant retina and littermate control, viewed by light microscopy (I) or electron microscopy (J). In controls, red blood cells (arrows) are located within vascular lumen. In mutants, extravascular red blood cells accumulate in NFL-GCL region. Asterisks in J: RGC somata. Abbreviations, see Fig. 3. Error bars: mean ± S.E.M. Sample sizes denoted by data points on graphs. Scale bars: 50 µm (A,C,E,G); 25 µm (I); 5 µm (J).

When microglia were ablated in this manner, astrocyte numbers failed to decline between P6 and P8, leading to an excess of ∼4,000 astrocytes relative to their littermate controls (Fig. 6C,D). This excess was not caused by increased proliferation (S3 Fig. I,J). After P8, astrocyte numbers began declining again in microglia-ablated animals, suggesting the existence of compensatory death mechanisms (see below for further investigation of this phenomenon). Despite this compensation, astrocyte numbers failed to return to normal levels by the end of the death period (Fig. 6D,F; S3 Fig. G).

To validate these results, we repeated the experiment using two different genetic strategies. First, we substituted a different microglial Cre line (a BAC transgenic *Cx3cr1-Cre* [54]) to drive DTR expression. Second, we examined *Csf1r* null mutant mice, which constitutively lack microglia [55]. In both cases, we confirmed that retinal microglia were absent before proceeding to assess astrocyte survival. Effects on astrocyte number were similar in magnitude to our original paradigm (Fig. 6E,F; S3 Fig. H). Together, these findings support the conclusion that microglia depletion impairs the developmental loss of astrocytes.

### Astrocytes that are spared in microglia-deficient mice survive and differentiate

The preservation of cells expressing astrocyte nuclear markers (Fig. 6C-F) could mean that microglia are required for astrocyte death; alternatively, this could mean that microglia are required merely for the clearance of dead astrocyte corpses, similar to their role in apoptosis [34]. In the former case, we would expect that the excess astrocytes in microglia-ablated mice should become mature, both molecularly and anatomically. However, if microglia mediate clearance but not death, ablation would be expected to increase the number of corpses without a similar increase in the number of differentiated astrocytes. To distinguish between these possibilities we stained microglia-ablated retinas for GFAP, a late marker of retinal astrocyte differentiation that also reveals cellular morphology [28,30,56]. Nearly all (>98.8%) Pax2^+^ or Sox9^+^ astrocytes co-expressed GFAP by P10, regardless of whether microglia had been ablated (Control: n=2,642 astrocytes; DTR: n=4,145 astrocytes; N=3 animals per condition). Thus, molecular differentiation of excess astrocytes was normal. Excess astrocytes also incorporated normally into the orderly mosaic of astrocyte cell bodies [30,57], suggesting that they engaged in normal repulsive interactions with their homotypic neighbors (S4 Fig. A,B). Further, the GFAP^+^ arbor network covered significantly more area in animals lacking microglia (Fig. 6G,H); the magnitude increase was similar to the increase in total astrocyte number (P10: 30% arbor area increase over littermate controls; 28% cell number increase). This finding suggests that excess astrocytes undergo morphological differentiation. Together, these results support the view that spared astrocytes are not corpses but rather have the capacity to develop normally.

### Retinal hemorrhage in the absence of microglia-mediated phagocytosis

We next investigated the functional role of microglia-mediated astrocyte death in retinal histogenesis. It has previously been shown that an excess of astrocytes leads to retinal hemorrhage, likely due to the close developmental association between astrocytes and vasculature [22,32]. We therefore hypothesized that a similar vascular phenotype should be evident in *Csf1r* mutant mice, which also have excess astrocytes (Fig. 6F). In *Csf1r* mutants, overall vascular patterning was largely normal, aside from some early subtle changes consistent with those previously reported for mice lacking microglia [58] (S4 Fig. E). However, histological examination of *Csf1r* mutant retina revealed bleeding within the RNFL. In thin plastic sections from P6 littermate controls (N=3), red blood cells (RBCs) were confined to the vasculature. By contrast, in mutants, numerous RBC profiles were observed throughout the RNFL extracellular space (Fig. 6I,J; S4 Fig. C,D; N=4, all were affected). Thus, microglia are required for vessel integrity during retinal angiogenesis. This finding is consistent with a model in which the microglial requirement for vascular integrity is mediated indirectly through microglial effects on astrocyte number.

### Compensatory mechanisms for astrocyte death in the absence of microglia

Ablation of microglia did not completely block astrocyte death; instead, death appeared to stop for ∼2 days before resuming again at virtually the same rate as in controls (Fig. 6D). A similar phenomenon was observed in *Csf1r* mutants: As astrocytes migrated centrifugally (Fig. 1A), their arrival in a given retinal region was followed by a ∼2 day period when astrocyte loss was impaired (S5 Fig. A). Subsequently, loss of astrocytes resumed with similar dynamics as controls, albeit with the decline curve shifted rightwards (S5 Fig. A,B). These observations suggest that accumulation of excess astrocytes triggers compensatory death when microglia are absent.

To investigate the nature of the compensatory mechanisms, we used *Csf1r* mutants to examine three possible alternate death routes. First, we tested whether astrocyte apoptosis might become the predominant death mechanism in the absence of microglia. At P6, the number of CC3^+^ astrocytes was higher in *Csf1r* mutants than littermate controls (Fig. 7A,B). However, this increase does not necessarily imply an increased rate of astrocyte apoptosis, because clearance of apoptotic corpses was also impaired in *Csf1r* mutants (Fig. 7D). Modeling studies indicated that failure of corpse clearance could entirely explain the increase in detectable CC3^+^ astrocytes, arguing against the possibility that more astrocytes become apoptotic in mutants (Fig. 7B). Further, CC3 staining in our acute DTR ablation model also failed to support the idea of elevated astrocyte apoptosis rates (S5 Fig. D). Thus, apoptosis is unlikely to be the compensatory astrocyte death mechanism.

**Figure 7.**
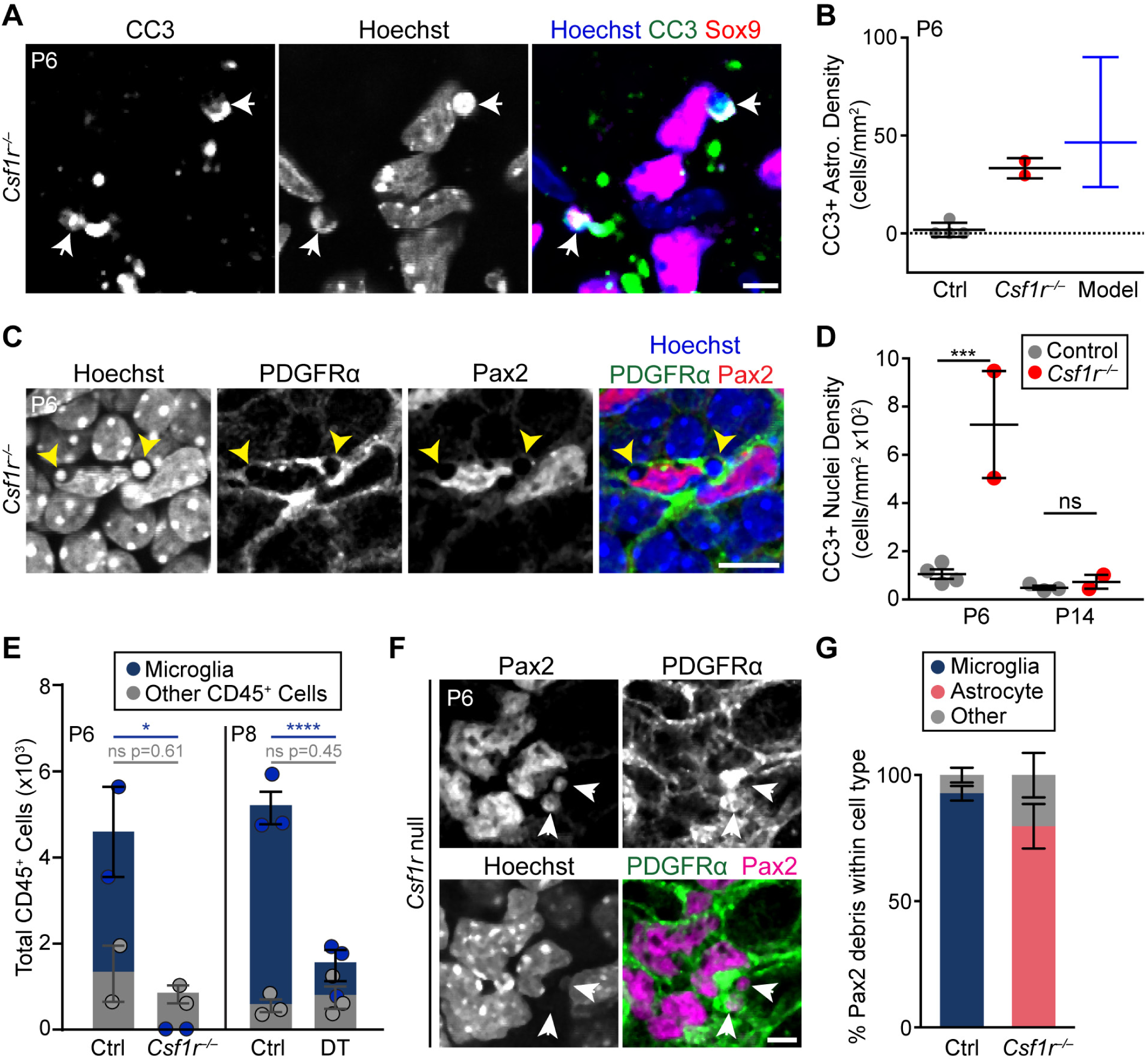
Compensatory mechaisms for astrocyte death in the absence of microglia. **A,B**) Example images (A) and quantification (B) of CC3^+^ astrocytes (arrows) in *Csf1r*^−/−^ retina. Model (B, blue) shows expected density of CC3^+^ astrocytes under a scenario where apoptosis occurs at a wild-type rate but no corpses are cleared (see Methods). Observed value from *Csf1r* null mutants (C, red) falls within 95% confidence interval of model. **C,D**) Non-professional phagocytes clear apoptotic corpses when microglia are absent. C: En-face view of *Csf1r*^*–/–*^ retinal whole-mount (single optical plane) showing pyknotic bodies (arrows, Hoechst channel) surrounded by PDGFRα^+^ astrocyte processes resembling phagocytic cups. Such features were never observed in wild-type mice. D: Delayed clearance of apoptotic corpses occurs between P6 and P14 in *Csf1r* mutants. All CC3^+^ nuclei within the NFL and GCL were quantified at P6 and P14. These are largely apoptotic neurons or endothelial cells; few are astrocytes (B). Statistics: 2-way ANOVA followed by Fisher’s LSD post-hoc test. P-values: P6, ***P=0.0006; P14, P=0.8340. **E**) Quantification of CD45^+^ cells in control and microglia-ablated retina. Non-microglial fraction of CD45^+^ cells (gray) was unchanged following acute (DT) or chronic (*Csf1r*) microglial ablation. Statistics: 2-way ANOVA followed by Fisher’s LSD post-hoc test. *p=0.0202; ****p<0.0001. **F**) En-face view (single optical plane) of RNFL in whole-mount *Csf1r* mutant retina. Pax2^+^ astrocyte debris localizes within PDGFRα^+^ astrocyte processes (arrows). **G**) Quantification of the percentage of Pax2^+^ debris localized to microglia, astrocytes or other cell types in control mice and in *Csf1r* mutants (P5-6). Most Pax2^+^ debris is found within astrocytes when microglia are absent. Error bars: mean ± S.E.M (except (C); blue: mean ± 95% confidence interval). Sample sizes denoted by data points on graphs. Scale bars: 5 µm (A,F); 10 µm (C).

Next, we tested whether loss of microglia causes another immune cell type to assume the role of phagocyte. *Csf1r*^*–/–*^ and littermate control retinas were stained with antibodies to CD45, a broad leukocyte marker. In controls, most retinal CD45^+^ cells were microglia, but a small population of non-microglial CD45^+^ cells (<2000 per retina) was present during the period of astrocyte death (Fig. 7E; S5 Fig. 5E). When microglia were eliminated using either *Csf1r* mutants or the toxin ablation model, the size of the remaining CD45^+^ population was similar to controls, arguing against any major infiltration of leukocytes (Fig. 7E; S5 Fig. E). Moreover, it is unlikely that the resident CD45^+^ cells were able to compensate on their own for the loss of microglia because we never observed any astrocyte material within these cells. For these reasons, leukocytes are poor candidates to mediate compensatory astrocyte elimination.

### Developing astrocytes engulf each other in the absence of microglia

Finally, we investigated a third possible route of compensatory astrocyte death: We asked whether microglia depletion might cause another retinal cell type to assume “death by phagocyte” functions. This is plausible because, in other contexts, non-professional phagocytes are able to take on engulfment responsibilities when professional phagocytes are absent [59,60]. Non-professional phagocytes appeared to be active in *Csf1r* mutant retina, since CC3^+^ apoptotic corpses were eventually removed following an initial clearance defect (Fig. 7D; S5 Fig. D). At least some of this compensatory corpse clearance was performed by RNFL astrocytes (Fig. 7C), raising the possibility that astrocytes might also take on other microglial phagocytic functions. We therefore tested whether absence of microglia might cause astrocytes to phagocytose each other. Using the Pax2 debris assay (Fig. 4A,C), we found that dead astrocyte material was readily identifiable in *Csf1r* mutants (Fig 7F). Unlike in controls, where such debris localized almost exclusively to microglia, debris in *Csf1r*^*–/–*^ retina was largely found within PDGFRα^+^ astrocytes (Fig. 7F,G; n=564 debris puncta from 3 controls, 93 ± 2.9% in microglia; n=111 debris puncta from 3 mutants; 80 ± 8.9% in astrocytes). This finding suggests that astrocytes themselves are the major cellular effector of astrocyte engulfment when microglia are absent. Our results are consistent with a model whereby astrocytes compensate for microglial ablation by killing and engulfing each other (Fig. 8).

**Figure 8.**
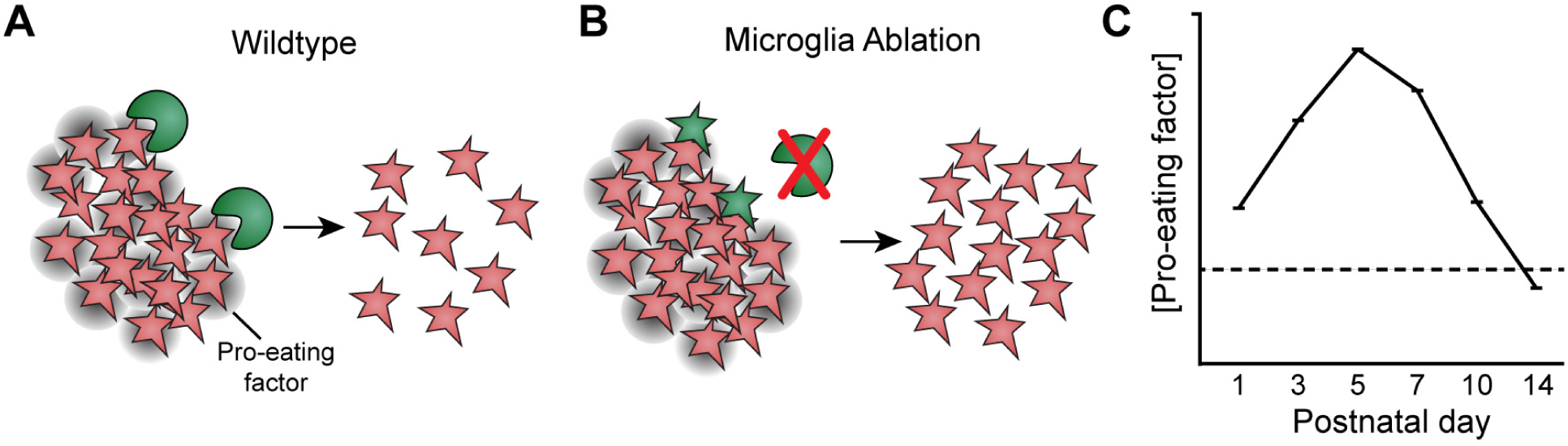
Model of the mechanisms driving astrocyte death. Schematic illustrating proposed mechanism of developmental astrocyte death, based on findings of this study. In wild-type mice (**A**), microglia (green) sense the number of astrocytes (red) within their local environment, perhaps due to expression of an astrocyte-derived pro-eating factor (gray shading; right panel). When astrocyte numbers are high, microglia are stimulated to remove them via phagocytosis. As astrocyte number falls, the amount of astrocyte-derived cue eventually becomes insufficient to stimulate microglia, thereby closing the death period (**C**). When microglia are ablated (**B**), death is temporarily halted. But astrocytes (green) eventually sense their own elevated numbers (red), leading them to phagocytose each other..

## Discussion

It has generally been assumed that developmental astrocyte death proceeds in a manner similar to neurons and oligodendrocytes, both of which die by apoptosis in large numbers [4,38]. However, these assumptions have rarely been tested in a quantitative manner; as a result, it remains unclear whether astrocytes are even subject to large-scale developmental death, let alone whether apoptosis is the mechanism. Here we demonstrate that retinal astrocytes undergo developmental death that is not apoptotic, but instead proceeds by a microglial “death by phagocyte” mechanism. The number of astrocytes killed by microglia is remarkably large: It includes not only the > 3-fold cell number decline between P5 and P14, but also a fraction of astrocytes that are consumed during their migratory period, prior to P5 (Fig. 3E; Fig. 4). Microglia are necessary for normal progression of astrocyte death, since death was reduced upon elimination of microglia from developing retina. Furthermore, we show that in the absence of microglia, astrocytes themselves become phagocytic and engulf their homotypic neighbors. This activity appears to partially compensate for the loss of microglia in mediating astrocyte removal. Nevertheless, elimination of microglia halts death for approximately two days, long enough to produce a ∼50% increase in the number of viable, differentiated astrocytes. This increase in astrocyte number may have functional consequences, since mice lacking microglia exhibit bleeding from the astrocyte-associated RNFL vasculature. Overall, these findings suggest that microglia-mediated death impacts the anatomy of the RNFL astrocyte network, and they raise the possibility that similar mechanisms may operate elsewhere in the CNS.

### Death by phagocyte as an astrocyte death mechanism

Five key findings support our conclusion that microglia are responsible for developmental astrocyte death. First, microglia in the RNFL assume a distinctive phagocytic phenotype, and consume astrocyte debris. Second, these behaviors are limited to the period when astrocytes are dying. Third, the frequency of observed astrocyte engulfment events is sufficient to explain the decline in astrocyte number. Fourth, experimental elimination of microglia reduces astrocyte loss. Fifth, the astrocytes preserved in microglial ablation experiments go on to differentiate, suggesting they have been spared from death. For these reasons we conclude that microglia kill retinal astrocytes. The phagocyte behavior we describe is fundamentally different from the well-described phenomenon of apoptotic corpse clearance, because interfering with phagocyte function does not typically rescue apoptotic cells from death [34] (S5 Fig. C). Astrocyte killing may be a case of “phagoptosis” – a form of cell death that can be prevented by blocking phagocytosis [14]. Since astrocytes may be engulfed while still viable (Fig. 4F), we favor the notion that phagocytosis is in fact the death-inducing event. However, our data do not rule out a scenario in which microglia first kill astrocytes via a separate (non-apoptotic) pathway before immediately engulfing the doomed cells. To accommodate this possibility, we have avoided the term phagoptosis and prefer the term death by phagocyte.

There is precedent for the notion that microglia can mediate death by phagocyte, both in pathological contexts [15–17] and during development of cortical subventricular zone progenitors [18]. Further, a recent study showed that retinal microglia can engulf embryonic RGCs [61], adding to the plausibility of our conclusions. One important difference between astrocytes and these other two cell types is that astrocytes rarely undergo apoptosis (Fig. 2). By contrast, RGCs and cortical progenitors are both subject to significant apoptotic death [39,42], suggesting that death by phagocyte may play more of a complementary role. Indeed, our results in *Bax* and *Csf1r* mutants indicate that apoptosis is a far more important determinant of RGC number than microglia-mediated death (Fig. 2B,D; S5 Fig. C). For astrocytes, by contrast, the opposite is true. Retinal astrocytes provide the first example, to our knowledge, of a CNS cell type that uses death by phagocyte as its primary mechanism of naturally-occurring developmental death.

Microglial killing of astrocytes is highly regulated – it occurs within a specific developmental epoch, and produces a relatively consistent population size by adulthood. What are the mechanisms that determine the timing and extent of astrocyte death? Our data suggest that microglia are driven to remove astrocytes due to signals derived from astrocytes themselves. A key indication that astrocytes produce pro-phagocytic signals is that microglia located in their vicinity (i.e. the RNFL) have a distinctive phagocytic phenotype. Moreover, several features of this phagocytic phenotype are expressed in a graded fashion that strongly correlates with the number of astrocytes present at any given age (see especially Fig. 3E but also Fig 3C,F,G; Fig. 4C,D). These findings are consistent with a model whereby microglia sense astrocyte number to modulate their phagocytic capacity (Fig. 8). In this model, the opening of the death period is driven by accumulation of excess astrocytes as they migrate into the retina, while its closing occurs when astrocyte numbers have dropped low enough to remove the phagocytic stimulus (Fig. 8C). Other factors, of course, could also contribute to the timing and extent of astrocyte death. For example, intrinsic developmental changes could influence the timing of when microglia are competent to remove astrocytes. The model we provide here (Fig. 8) should serve as a useful framework for exploring such possibilities, and thereby dissecting the cellular and molecular mechanisms that regulate astrocyte population size.

### Compensatory astrocyte-mediated astrocyte death in the absence of microglia

When professional phagocytes are unavailable, non-professional phagocytes carry out engulfment functions such as clearing apoptotic corpses [59,60]. Here we show that retinal astrocytes can substitute for microglia not only in this capacity, but also in the engulfment of their astrocyte neighbors (Fig. 7F,G). This activity is likely responsible for resumed astrocyte death following microglia ablation, as we did not find evidence for other potential death routes. Müller glial cells (or their radial glial progenitors) may also serve as non-professional phagocytes in other contexts [62], but their contribution here appears minor because ∼80% of astrocyte debris was localized within other astrocytes.

What causes astrocytes to begin engulfing each other when microglia are absent? In the case of apoptotic corpse clearance, a key feature of the compensatory mechanism is that both microglia and non-professional phagocytes have intrinsic capacity to recognize and clear dying cells. However, non-professional phagocytes are far slower to begin engulfing an apoptotic cell after first encountering it [59]. This delay reserves non-professional phagocytosis for instances where microglia cannot keep up. Once engulfment has begun, however, both cell types clear dead cells at similar rates [59]. For retinal astrocytes, the timecourse of compensatory death (Figs. 6D; S5 Fig. A,B) suggests that precisely this mechanism could be at play. We show that the rightward shifts of these curves are due almost entirely to a delay in onset of cell loss; once loss begins, the dynamics of the decline phase are quite similar between wildtype and ablated retinas (S5 Fig. B). These data are well explained by a model in which astrocytes are slow to begin engulfing each other when the need first arises, but once removal is underway, it proceeds with a mostly normal timecourse. In this model, astrocytes, like microglia, are sensitive to pro-phagocytic astrocyte-derived cues (Fig. 8), but they normally do not have time to act on these cues because microglia do so first. When microglia are absent, the latent propensity for developing astrocytes to engulf each other is unveiled.

### Relevance to astrocyte death in other CNS regions

Few previous studies have addressed quantitatively the extent of astrocyte death in the developing CNS. It was not known if astrocytes, like many types of neurons, are overproduced and then culled. Here we show that at least one CNS astrocyte population follows such a pattern. Studies of absolute astrocyte number will be needed to see if this holds true for other CNS regions.

Extrapolating from apoptosis assays, it was reported that astrocytes in cortex [23,25,26] and retina [27–29] were subject to developmental death. However, quantitative information about the rates of astrocyte apoptosis, or its impact on total cell numbers, was not included in these studies, leaving open the possibility that apoptosis has only a minor effect on population size. Indeed, in several of these studies, astrocyte apoptosis was remarkably infrequent [23,26]. A study in rat cerebellum showed that a large fraction – as many as 1% – of GFAP^+^ white matter cells are apoptotic at P7 [24]. At the time of this study it was thought that these GFAP^+^ cells were astrocytes, but it is now known that they are actually multipotent neural progenitors rather than lineage-committed astrocytes [63]. Thus, astrocyte apoptosis rates in the cerebellum are likely much lower than originally proposed. Our retinal data indicate that microglia-mediated death is a far more significant contributor to astrocyte loss than apoptosis. It will be interesting to learn whether microglia also kill astrocytes in other regions of the CNS. As in the RNFL (Fig. 3F,G), a transient population of Osteopontin^+^ phagocytic microglia is found in developing brain [46,64], raising the possibility that astrocytes could be targets of such cells. In this regard, it is noteworthy that microglia ablation increases expression levels of the astrocytic protein ALDH1L1 in brain [43]. Whether this reflects increased brain astrocyte numbers remains to be determined.

### Potential functions of astrocyte death in retinal development

Retinal astrocytes play a pivotal role in retinal angiogenesis, guiding vascular colonization of the RNFL during the perinatal period (P0 until ∼P8 in mice; Fig. 1A; [22]). This guidance is accomplished both through direct physical patterning of growing vessels, as well as expression of key pro-angiogenic signals such as VEGF [22,30,65]. Changes in the size of the astrocyte population would be expected to disturb angiogenesis, by altering VEGF levels and by changing the pattern of the astrocyte network in a manner that could be propagated to vessels. Accordingly, when astrocyte numbers are artificially elevated through forced proliferation, patterning of the astrocyte and vascular networks are both perturbed. Crucially, these perturbations cause retinal bleeding during the early postnatal period [32]. This result highlights the importance of developmental mechanisms that maintain appropriate astrocyte numbers, and the deleterious consequences when such mechanisms are dysregulated.

Here we show that astrocytes are subject to developmental cell death during the period of RNFL angiogenesis. We propose that death may be part of a homeostatic mechanism that balances migration and proliferation to set the astrocyte population size. In this case, both increased proliferation [32] and decreased death (this study) would be expected to have similar effects. In *Csf1r* mutants, the magnitude of effects on the astrocyte network were smaller than in the proliferation study [32], likely due to the fact that we only partially blocked death. However, in both cases the number of astrocytes and the density of their arbor network was increased. Furthermore, in both cases retinal bleeding was observed during early postnatal development (Fig. 6I,J; [32]). The many phenotypic similarities between these two mouse models suggest that bleeding in *Csf1r* mutants might be astrocytic in origin. In our study we cannot exclude that bleeding was caused by an astrocyte-independent function of microglia, but in the Fruttiger et al. (1996) study the manipulation was likely astrocyte-specific. The presence of excess VEGF-producing astrocytes is a plausible cause of hemorrhage, since VEGF tends to promote endothelial cell proliferation and sprouting at the expense of quiescence, maturation, and barrier formation [66]. In order to stringently test the role of death in vascular development, it will be necessary to identify molecular manipulations that prevent microglia from killing astrocytes without eliminating microglia entirely. Despite extensive efforts (Fig. 5), we were unable to identify such a manipulation in this study. However, we expect that this will ultimately become possible, thereby enabling definitive studies on the functional role of developmental astrocyte death.

## Materials and Methods

### Experimental Model and Subject Details

#### Animals

The Institutional Animal Care and Use Committees at Duke University and the University of California at Santa Barbara reviewed and approved all experimental procedures involving animals at respective universities. Mice were maintained on a 12 hour light/dark cycle; food and water were freely and continuously available. All mouse strains used (see Key Resources table) were maintained by continual backcrossing to C57Bl6/J, unless noted below. Upon receipt into our colony, each strain was genotyped for *Rd1* and *Rd8* retinal degeneration mutations; where necessary, these mutant alleles were bred out of the strain. As such, all animals used for experiments were free of these retinal degeneration mutations. For Cre-dependent reporter expression, we used both the *Rosa26*^*ai14*^ allele [69], which drives tdTomato expression, and *Rosa26*^*iDTR*^ allele, which drives cell-surface expression of the human DTR; in the latter case, anatomy was assessed by antibody staining to DTR (see Method Details below). *Cx3cr1*^*CreER-ires-YFP*^ mice ([43]; also denoted *Cx3cr1*^*CreER*^) served three different purposes in this study: 1) microglia-specific Cre driver; 2) constitutive microglia-specific YFP expression; 3) *Cx3cr1* loss-of-function allele (Fig. 5).

To generate *Csf1r* null mutants (*Csf1r*^*–/–*^), *Actin-cre* animals were crossed to *Csf1r*^*flox*^ animals to allow for germline recombination of *Csf1r*. As *Csf1r* null mutations are lethal at birth on a pure C57Bl6/J background (data not shown), we used a hybrid C57Bl6-SJL strategy to generate viable *Csf1r* null mutants. In this strategy, two sets of breeders were used for experimental matings. One set (denoted B6-*Csf1r*^*+/–*^) was produced by continual backcrossing of the *Csf1r* null allele to C57Bl6/J; the other set was produced by outcrossing the B6-*Csf1r*^*+/–*^ mice to SJL/J, thereby generating F1 hybrids of B6 and SJL that carried the *Csf1r* null allele (denoted B6SJL-*Csf1r*^*+/–*^). Experimental litters were generated by crossing B6-*Csf1r*^*+/–*^ to B6SJL-*Csf1r*^*+/–*^. Mutant progeny routinely survived to ∼P21 without special husbandry. Note that, due to this special breeding strategy, it is exceedingly difficult to deploy Cre-lox reporters in the *Csf1r* background: Since Cre and reporter lines are not available on an SJL background, crossing them into the *Csf1r* background would likely result in neonatal lethality.

For all experiments, controls were *Csf1r*^+/+^ littermates and at least one eye per animal was stained for microglia to confirm their absence.

#### Tamoxifen administration

To induce CreER-mediated recombination, 100 μg of tamoxifen, an estrogen receptor ligand, was administered via intraperitoneal (i.p.) injection to *Cx3cr1*^*CreER*^ neonatal mice. Tamoxifen powder was dissolved in corn oil at a concentration of 20 mg/mL by sonicating in a room temperature water bath for 30 min. Injections were administered at P2, P5, and P7 for DTR experiments, and at P0 or P1 for *Syk*^*flox/flox*^ experiments. Both control and experimental animals received tamoxifen injections in all experiments that required CreER-mediated recombination, unless otherwise noted.

#### Diphtheria toxin-mediated microglia ablation

For microglia ablation experiments, the *Rosa26*^*iDTR*^ line [70] was crossed to two different microglial Cre drivers: 1) *Cx3cr1*^*CreER-ires-YFP*^ (see above); and 2) a *Cx3cr1-Cre* BAC transgenic line [54]. Most experiments were performed on the *Cx3cr1*^*CreER*^ background. To establish an ablation paradigm for neonatal mice, we first tested the specificity, efficacy, and temporal characteristics of the *Cx3cr1*^*CreER*^; *Rosa26*^*iDTR*^ ablation paradigm in neonatal retina. Administration of tamoxifen at P2 induced cell-type-specific expression of DTR in nearly all microglia by P4 (S3 Fig. A; 97.41 ± 0.52% of microglia were DTR^+^; 100% of DTR^+^ cells were microglia; n = 580 cells from 2 mice, mean ± S.E.M.). DTR-expressing mice, but not littermate controls lacking either the CreER or the DTR transgenes, were susceptible to microglia ablation upon administration of diphtheria toxin: A single 80ng dose depleted microglia to ∼10% of control levels within 2 days (Fig. 6B; S3 Fig. B). Depletion was confirmed by staining for 3 different microglial markers (Iba-1, CD45, and YFP driven by the *Cx3cr1*^*CreER*^ allele; S3 Fig.E; Fig. 7E; S5 Fig. E). Microglia were specifically affected – we did not observe any effects of diphtheria toxin on overall retinal histology (S3 Fig. D; [30]). By 4 days post-injection, however, microglia had partially repopulated the retina, returning to ∼50% of control levels, only ∼60% of which expressed DTR (S3 Fig. B,C).

To ablate microglia for longer periods of time, we developed a regime for repeated tamoxifen and DT administration; this successfully prevented return of microglia for the entire administration period – as long as P6-14 (Fig. 6A,B). In this regime, tamoxifen was administered as described above to induce DTR expression; diphtheria toxin (DT) was then administered at P4, P6, P8, P10, and P12 to allow for continuous depletion of microglia. DT, dissolved in 1x PBS, was administered i.p. at 80ng per dose; we previously found this dose to be optimal for ablation without off-target effects in neonatal pups [30]. For most experiments, the control group was littermates that were injected with both tamoxifen and DT but did not inherit one of the two key transgenes (i.e. *Cx3cr1*^*CreER*^ or *Rosa26*^*iDTR*^). In initial experiments we compared control animals of this type to those that received only tamoxifen but no DT (or, in two cases, animals that received neither; these animals are denoted by gray dots with black outlines in S3 Fig. F). There was no difference between these control groups in total astrocyte number counts at P8 and P10 (S3 Fig. F), so data from both control groups were ultimately combined for final analyses (Fig. 6; S3 Fig.). For DTR-mediated microglia ablation experiments in the *Cx3cr1-Cre* background, the same DT dosage described above was administered to both control and experimental animals at P6 and P8. Control animals were littermates that lacked either the *Cx-3cr1-Cre* or *Rosa26*^*iDTR*^ transgenes.

#### Minocycline Administration

For minocycline experiments, CD-1 mice were i.p. injected once daily with minocycline (Sigma; 50mg/kg) or vehicle (10mM Tris-HCl) from P4-P9. Dosage was determined based on previously published studies [71].

### Histology

Mice were typically anesthetized and euthanized by decapitation, followed by immediate enucleation. Whole eyes were fixed on ice in 4% paraformaldehyde (PFA) in 1X PBS for between 1.5 and 2 hours, and subsequently washed twice (5 minutes each) in 1X PBS. In some cases, mice were perfused transcardially with 4% PFA/1X PBS prior to eyeball post-fixation as above. Eyes were either processed for histology immediately following fixation or stored at 4°C in 0.02% Sodium Azide in 1X PBS until the time of tissue processing. Retinas were prepared for cryosections or whole-mounts as previously described [72]. Briefly, for retinal sections, the lens was extracted from the eyecup and vitreous removed. Eyecups were then cryoprotected in 30% sucrose for at least 2 hours prior to embedding in Tissue Freezing Medium. After freezing, 20 µm cryosections were collected with a Microm HM 550 cryostat. For whole-eye sections, vitreous structures were maintained by leaving both retina and lens within the eye cup and puncturing the cornea with a 30 Gauge needle followed by a slight elongation of the puncture with iris scissors to allow for thorough cryoprotection. After cryoprotection, whole-eyes were embedded in Tissue Freezing Medium and 20 µm cryo-sections were obtained as previously described. Whole-mount retinas were obtained by also extracting lens and vitreous, in addition to detaching the retina from the eyecup.

### Immunohistochemistry

Retinal tissue was blocked in a solution of 3% normal donkey serum, 0.3% Triton-X and 0.02% Sodium Azide in 1X PBS at room temperature (30 minutes for retinal sections; 2 hours for wholemount retina). After blocking, tissue was incubated with primary antibodies in the aforementioned blocking solution either overnight (retinal sections) or for 5 days (wholemount retina). Tissue was washed after primary antibody incubation at least three times in 1X PBS. Secondary antibodies and Hoechst 33258, in a solution of 0.3% Triton-X in 1X PBS, were then applied for either 2 hours at room temperature (retinal sections) or overnight at 4°C (wholemount retinas). Finally, tissue was washed at least three times at room temperature before preparing samples for image acquisition.

### Image acquisition and processing

Following immunohistochemistry, whole retinas were placed in a dish of 1X PBS and four radial cuts separated by 90 degrees were made, with each cut extending approximately 1/3^rd^ of the way from the edge of the retina to the optic nerve head. Cut retinas were placed ganglion cell-side up on nitrocellulose filter paper and carefully laid flat with a paintbrush before mounting on a slide. Prior to imaging, slides were coverslipped with a layer of Fluoromount G mounting media between the sample and coverslip. The coverslip was held in place with a layer of clear nail polish applied at the seam between the edge of the coverslip and the slide. A Nikon A1 or an Olympus FV300 confocal microscope was used to image sections and whole-mounts. To produce large field-of-view images, tiled frames were acquired with a 20x air objective via the Nikon A1 confocal or an Olympus IX81 epifluorescence microscope, using automated software supplied by the microscope manufacturer. Images were then stitched into a single image using Olympus or Nikon software. For section and whole-mount images, z-stacks were collected at a z-resolution of 0.3–1 µm; 1.5 µm z-resolution was used for tilescan z-stacks. Image stacks were imported to Fiji [73] for processing and analysis. Images selected for display were first maximum projected to a single plane prior to de-noising by median-filtering (2.0 pixel radius for most images; in rare instances pixel radius >2.0 and up to 10.0 was utilized). The portion of the stack selected for maximum-intensity projection was determined by the z-volume of the structure to be depicted in the final image. Assembly of color channels as well as minor adjustments to brightness and contrast were also made in Fiji. For data analysis and quantification, only original stacks and not z-projections were used, unless otherwise noted.

### Plastic Sections and Electron Microscopy

Immediately following enucleation, whole eyes were fixed in 2% PFA + 2% glutaraldehyde in 1X PBS for 1 hour at room temperature, and then overnight at 4°C. Eyeballs were then washed twice in 1X PBS and immersed in 2% osmium tetroxide in 0.1% cacodylate buffer, dehydrated, and embedded in Epon 812 resin. For light microscopy, semi-thin sections of 0.5 µm were prepared and counterstained with 1% methylene blue. For electron microscopy, thin sections of 65-75nm thickness were collected on a Leica EM CU7 and counterstained with a solution of 2% uranyl acetate and 3.5% lead citrate, and examined using a JEM-1400 transmission electron microscope at 60 kV. An Orius 1000 charge-coupled device camera was used to collect images. Astrocyte nuclei were identified by their localization proximal to RGC nerve fibers, the oblong shape and size of their nucleus, and their unique heterochromatin pattern. As pericytes also have a nuclear morphology similar to that of astrocytes, we distinguished between these cell types based on the characteristic tight association between pericytes and endothelial cells. Red blood cells were identified by their characteristic irregular shape and electron-dense staining profile.

### Modeling

#### Overall strategy

To estimate the contribution of apoptosis and phagocytosis to developmental astrocyte loss, a model originally formulated to estimate developmental cortical neuron death was applied to our system [35]. This model predicts the number of cells that remain at the end of the death process (*N*_*T*_) based on the rate at which cells are dying and the starting number of cells (*S*). The rate of death is determined by two parameters: 1) The fraction of the cell population that is visibly dead or dying on a given day (*D*_*n*_), and 2) The amount of time for which a dead cell is visible on day *n* (*V*_*N*_; this can also be thought of as a clearance time). By assuming that *D*_*n*_ and *V*_*n*_ are constant (i.e. reflecting an average across all days of development), we were able to simplify the model to the following equation:

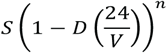

If the starting number and final number of cells as well as the fraction of dead cells in a given death process are known, clearance time can be determined by solving for *V*:

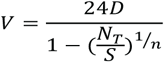

To validate that this model can accurately estimate cell death under the conditions of our assumptions, we utilized it to predict *V* (visibility, or clearance time) for RGCs, as this is a cell population known to undergo apoptosis during normal development [37]. We used previously published rat RGC developmental data [36] to carry out this validation. Note that in the original publication, data for RGC counts and total number of degenerating neurons in the ganglion cell layer (GCL) were listed with ages 0 to 10 days after birth. These ages are related to the date of injection for RGC retrograde labeling with Horseradish Peroxidase (HRP), while actual RGC quantification was done 18 hours post-injection – therefore, day 0 is actually P1, day 3 is actually P4, and so on. Thus, for our purposes, we are calling “0 to 10 days after birth” now “P1 to P11.” Based on this paper [36], the following parameter values were used for the model (also see S2 Fig. D): *D* was the average percentage of neurons in the GCL found to be pyknotic between P1 and P11 (*D* = 0.005375, or 0.5375%); *S* = 200,000 RGCs (P1); *N*_*T*_ = 117,000 RGCs (P11); and *n* = 10 days (P1 to P11). While *D* in this case is not specific to RGCs (given that amacrine cells also reside in the GCL), we made the assumption that all neurons of the GCL undergo a similar rate of degeneration and thus treat *D* as if it were the rate of RGC death. Based on the fact that our predicted *V* for RGCs (Fig. 2B) was in good alignment with clearance rate measurements made independently by others [37–40], we concluded that the assumptions underlying our model were sound. As such, we proceeded to use the model to investigate astrocyte death.

#### Modeling apoptosis as an astrocyte death mechanism

To estimate the contributions of apoptosis to astrocyte death, we denoted to be the average number of astrocytes found at P5 (thus, *S* = 26,987 cells). This age was chosen for several reasons. First, P5 is when astrocyte numbers are at their highest. Additionally, astrocytes have finished migratory colonization of the retina by this age, eliminating the need to account for cellular migration when predicting the relationship between total astrocyte numbers and the contributions of cell death. *D* was the experimentally determined average percentage of CC3^+^ astrocytes during the P5 and P14 period (thus, *D* = 0.000583, or 0.0583%, see S2 Fig. C). *n* was set to 9 (i.e. the length of the P5-P14 death period). These three parameters were held constant for all three analyses described below.

We used the model to address three different questions related to astrocyte apoptosis. First, we asked what value of clearance time *V* would be required to account for the change in astrocyte number between P5 () and P14 () if the death rate were the observed *D.* For this calculation, *N*_*T*_ was the average number of astrocytes found at P14 (thus, *N*_*T*_ = 6,513 cells). The computed value of *V* (Fig. 2C, blue curve) was on the order of ∼5 minutes. In a second permutation of the model, we asked whether the observed death rate *D* could predict the observed if the clearance time *V* were set to a biologically plausible value. For this purpose we used the clearance time for apoptotic RGCs (2.47 hours) as *V.* The outcome of this calculation is the red curve plotted in Fig. 2C.

Third, we used the model to estimate the total number of astrocytes that died by apoptosis in wild-type mice. This number was then used to estimate the expected CC3^+^ astrocyte density at P6 in a situation where apoptotic corpses could not be cleared (e.g. in mice lacking microglia). To make the total number estimate, the same model parameters were used as in the second scenario above to generate an astrocyte numbers timecourse (e.g. Fig. 2C, red line). The number of astrocytes lost each day was determined from the difference between astrocyte numbers on day N and day N+1 for each pair of days. We then calculated the average number of astrocytes lost per day and multiplied this average by six days to arrive at an estimate of astrocytes lost to apoptosis between birth and P6. To put confidence bounds on this estimate, we determined the 95% confidence interval of our CC3^+^ astrocyte number measurement (S2 Fig. C), and performed the same analysis again using the upper and lower confidence interval values for the death rate *D* parameter. Thus, the calculation of “astrocytes lost” between P0 and P6 was ultimately performed three times with three different *D* values (*D*_*mean*_ = 0.03%; *D*_*min*_ = 0.02%; *D*_*max*_ = 0.06%), generating a “mean” as well as an upper and lower limit to our estimate of astrocytes lost. Finally, we used these values to calculate the expected density of CC3^+^ astrocytes at P6 if none of the apoptotic astrocytes that died prior to P6 were ever cleared (Fig. 7B, blue line). The conversion from total number to density was made by dividing by the average retinal area at P6.

#### Modeling phagocytosis as an astrocyte death mechanism

To calculate *V* as it relates to the contributions of phagocytosis to astrocyte death, we again denoted *S* to be the average number of astrocytes found at P5 (thus, *S* = 26,987 cells). *D* was the experimentally-determined average percentage of astrocytes found to be fully enveloped by microglial processes during the P5-P14 period (thus, *D* = 0.00556, or 0.556%). As before, *N*_*T*_ was the average number of astrocytes found at P14 (thus, *N*_*T*_ = 6,513 cells) and *n* was 9 (P5 to P14).

### Quantification and Statistical Analysis

#### Statistics

Error bars are expressed as mean ± S.E.M., unless otherwise noted. For all analyses, alpha was set to 0.05. Statistical parameters (i.e. sample size, statistical and post hoc tests, and statistical significance) are reported in every figure or figure legend. All t-tests were two-tailed. Two-way or Three-way ANOVAs without matching, followed by Uncorrected Fisher’s LSD or Tukey’s multiple comparisons tests, respectively, were utilized when appropriate. Asterisks denote statistical significance (*, p ≤ 0.05; **, p ≤ 0.01; ***, p ≤ 0.001; ****, p ≤ 0.0001) in every figure. Data were analyzed using GraphPad Prism v7 software.

#### Quantification of Astrocyte Density and Total Astrocyte Number

We sampled astrocyte density from central, middle, and peripheral regions of the retina (Fig. 1B). Typically, 3-4 60x images per retinal region were acquired for the analysis, although in rare cases only 2 images were available. Only retinal locations free of damage were sampled. These locations were chosen using anatomical features of the retina so as to maintain consistency in sample selection. Central images were chosen by sampling retinal tissue just proximal to where blood vessels first bifurcate upon exiting the optic nerve head into the retina. Middle images were acquired at the approximate midpoint of the line connecting the edge of the retina to the optic nerve head. Finally, peripheral images were acquired ∼2 to 3 astrocyte cell bodies away from the edge of the retina. Astrocyte density for each image was quantified by hand using ImageJ/ FIJI software to mark each cell. The average density per region was then converted into an overall weighted average density based on the total retinal area within each sampled region (weighting: central=11%, middle=33%, peripheral=56%; see Fig. 1B and [30]). This weighted average was multiplied by retina area to obtain a value for estimated total astrocyte number. Retina areas were measured in FIJI by drawing perimeters with the Freehand selection tool on tilescanned images of wholemount retinas.

#### Microglia-Astrocyte Interaction Analysis

Images were collected from retinal whole-mounts by sampling central, middle, and peripheral retina as described above. Microglial morphology was revealed in *CX3CR1*^*CreER-ires-YFP*^ mice using anti-GFP immunohistochemistry. To quantify whether an astrocyte was touched, partially enveloped or fully enveloped by a microglial process, we focused on such interactions occurring only at astrocyte somata labeled with either Pax2 or Sox9. Any astrocyte soma found to be contacted by a microglial process was labeled as “touched.” If an astrocyte soma was found to have at least 50% of its circumference enveloped by a microglial process, it was labeled as “partially enveloped.” Astrocyte somata were counted as “fully enveloped” if the entire circumference of the astrocyte soma was surrounded by a microglial process.

#### Lysosome Index

To quantify microglial lysosomal content, 60X confocal z-stacks which captured the entire depth of the RNFL plus GCL (∼12 µm total, 1µm z-steps) were obtained from retinal whole-mounts stained for Iba1 (to identify microglia) and CD68 (to identify their lysosomes). Sampling methodology from central, middle, and peripheral retina was as described above. Laser settings were individually adjusted for each image to ensure under-and over-exposure of CD68^+^ pixels did not occur. Stacks were processed into maximum intensity z-projections using Fiji software. On rare occasions, there were images which contained CD68^+^ endothelial cells – such images were discarded in this analysis. In Fiji, CD68^+^ pixels were segmented using thresholding (settings: Li, black and white, dark background). The ImageJ function “Analyze Particles” was used to determine the percentage of image area occupied by CD68^+^ pixels (Analyze Particles settings: size 0-Infinity, analyzed in pixel units, circularity 0.00-1.00, include holes). The number of microglia within the given image was quantified, and the number of CD68^+^ pixels normalized to the number of microglia to obtain a Lysosome Index for the given image. For cross-developmental comparison between Lysosome Index and total astrocyte number, both measures were normalized to maximum values and plotted on a 0-100 scale, using GraphPad Prism software.

#### Debris Quantification

Pax2^+^ or tdTomato^+^ astrocyte debris were classified as engulfed by microglia only if the debris were found to be enclosed on all sides by microglia or their phagosomes. This was determined by examining confocal z-stacks encompassing the entire microglial cell (z step 0.4-1.0 µm). A similar analysis was performed to assess astrocytic engulfment of Pax2^+^ debris in *Csf1r* mutants. For wild-type Pax2^+^ debris studies, microglia were labeled by staining for anti-GFP in *Cx3cr1*^*CreER-ires-YFP*^ mice (P3, P5, P10, and P14). For *Csf1r* mutant studies, the strain was as described in “Animals” section above (P6). Note that different lots of the Pax2 antibody supplied since 2015 have exhibited higher background, making them unsuitable for debris analysis. In the lots we received prior to that time, background was extremely low, making identification of debris quite simple. We conducted follow-up studies using tdTomato in order to rule out the possibility that the debris was an artifact of the original Pax2 antibody lots.

#### Astrocyte Network Analysis

Images were collected from GFAP-stained retinal whole-mounts by sampling central, middle, and peripheral retina as described above. For quantification of astrocyte network coverage, 60X confocal z-stacks which captured the entire depth of the RNFL plus GCL (∼12 µm total, 1µm z-steps) were processed into maximum intensity z-projections using Fiji software. Only central and middle retina were sampled as GFAP expression is low in peripheral retina at P10. Total network coverage was quantified by measuring the percentage of GFAP negative space within a given ROI, and subtracting this percentage from 100 to obtain the percentage of space occupied by the GFAP astrocyte network. GFAP negative space was segmented using automatic thresholding (settings: Triangle), followed by application of the ImageJ function “Analyze Particles” (Analyze Particles settings: size 0-Infinity, analyzed in pixel units, circularity 0.00-1.00). Data was normalized to control by litter prior to statistical analysis due to litter-to-litter variability.

For quantification of astrocyte network regularity, 60x confocal z-stacks through the RNFL and GCL were acquired from P10 retinas stained with antibodies to GFAP and Pax2 or Sox9. Using Fiji software, a dot was manually placed at the center of each cell to generate X–Y coordinates. These coordinates were used to produce Voronoi domain regularity indices using Fiji software, as previously described [74].

### Data and Software Availability

All materials, datasets, and protocols used in this manuscript are available from the corresponding author on reasonable request. Software utilized in the current study are listed in the Key Resources Table.

## Key Resources Table

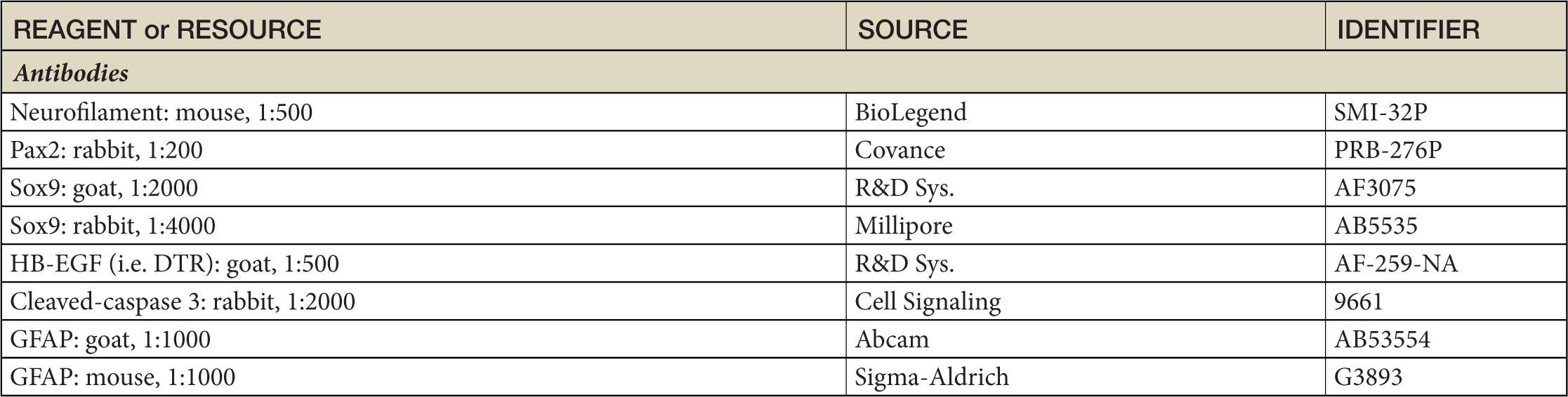
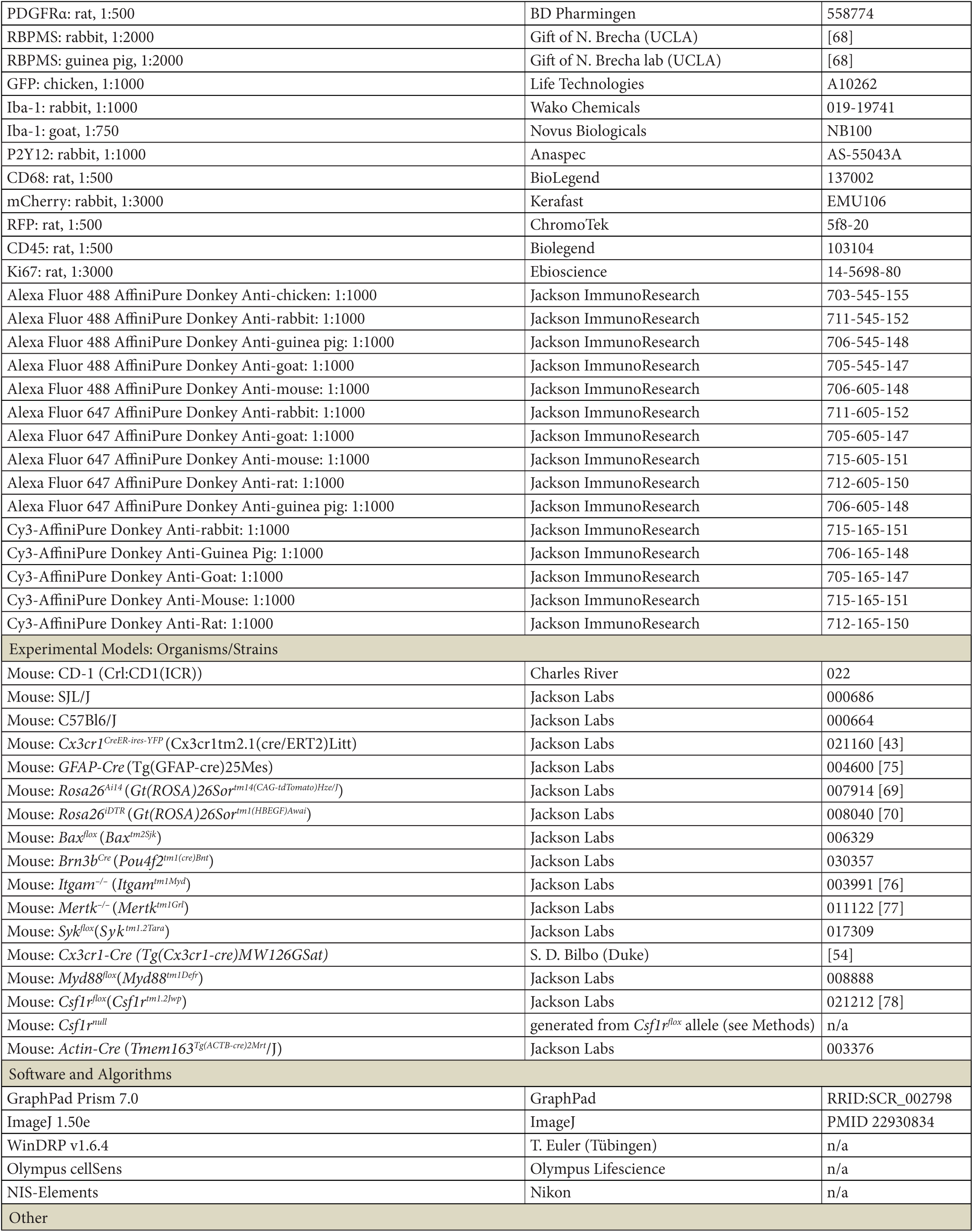
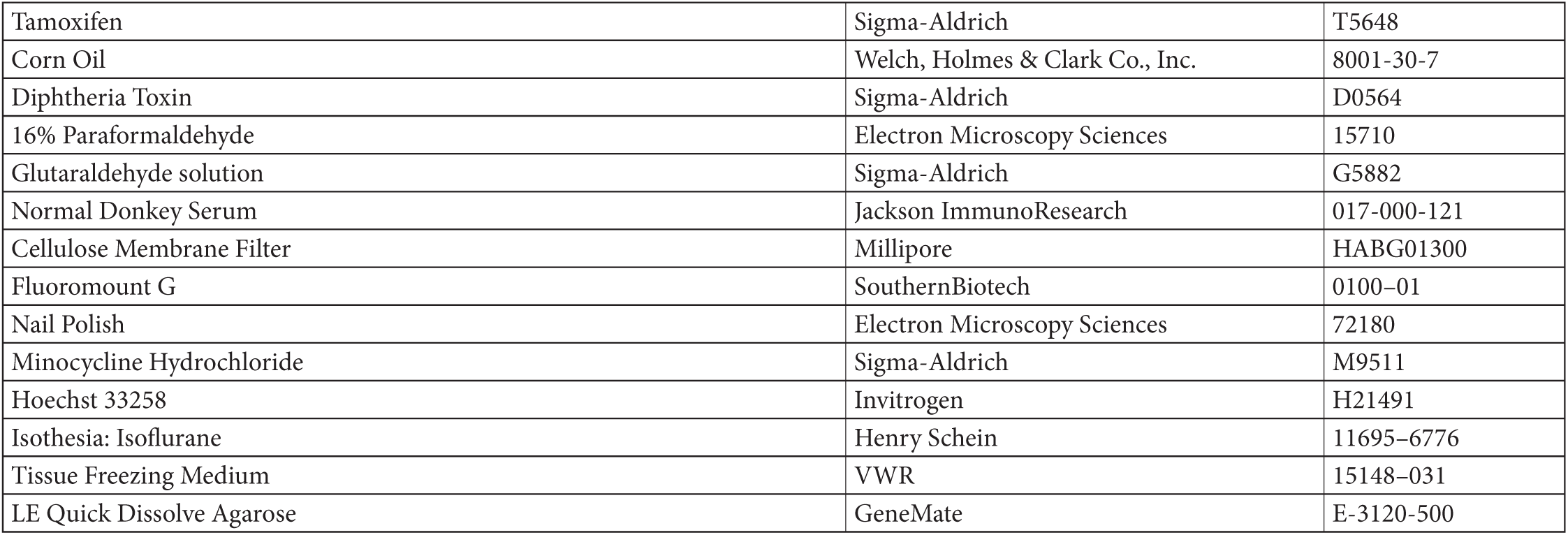

### Acknowledgments

The work was supported by grants from the National Institutes of Health (EY024694 to J.N.K; EY019968 to B.E.R.; EY5722 to Duke University); Ruth K. Broad Foundation to J.N.K.; Duke University Holland-Trice award to J.N.K.; McKnight Endowment Fund for Neuroscience to J.N.K.; Pew Scholars award to J.N.K.; National Science Foundation Graduate Student Research Fellowship (DGE-1644868) to V.M.P.; Duke Neuroscience Summer Program of Research to F.S.B.; and a Research to Prevent Blindness Unrestricted Grant to Duke University. We thank Ari Pereira for mouse husbandry; Claire Yin, Varun Pai, and Jaesook Yoo for technical assistance; Ying Hao for assistance with histology and electron microscopy; Susan Ackerman (UCSD) for suggesting the SJL hybrid vigor breeding strategy; Staci Bilbo for providing the *Cx3cr1-Cre*; *MyD88*^*flox*^ breeders; Greg Lemke (Salk) for providing *Mertk* mutant eyes; Nicholas Brecha (UCLA) for providing RBPMS antibodies; and Cagla Eroglu for comments on the manuscript. The funders had no role in study design, data collection and analysis, decision to publish, or preparation of the manuscript.

## Author Contributions

Conceptualization: VMP, CEP, FSB, JNK. Methodology: VMP, CEP, FSB, MAL, ERO, DRS, RMP, CRA, BER, JNK. Formal analysis: VMP, CEP, JNK. Investigation: VMP, CEP, FSB, MAL, RMP, JNK. Resources: CRA, BER. Writing – Original Draft: VMP, JNK. Writing – Review & Editing: VMP, CEP, FSB, MAL, RMP, ERO, DRS, CRA, BER, JNK. Visualization: VMP, JNK. Funding Acquisition: VMP, JNK.

## Supplementary Data

### Supplementary Figures

**Figure S1:**
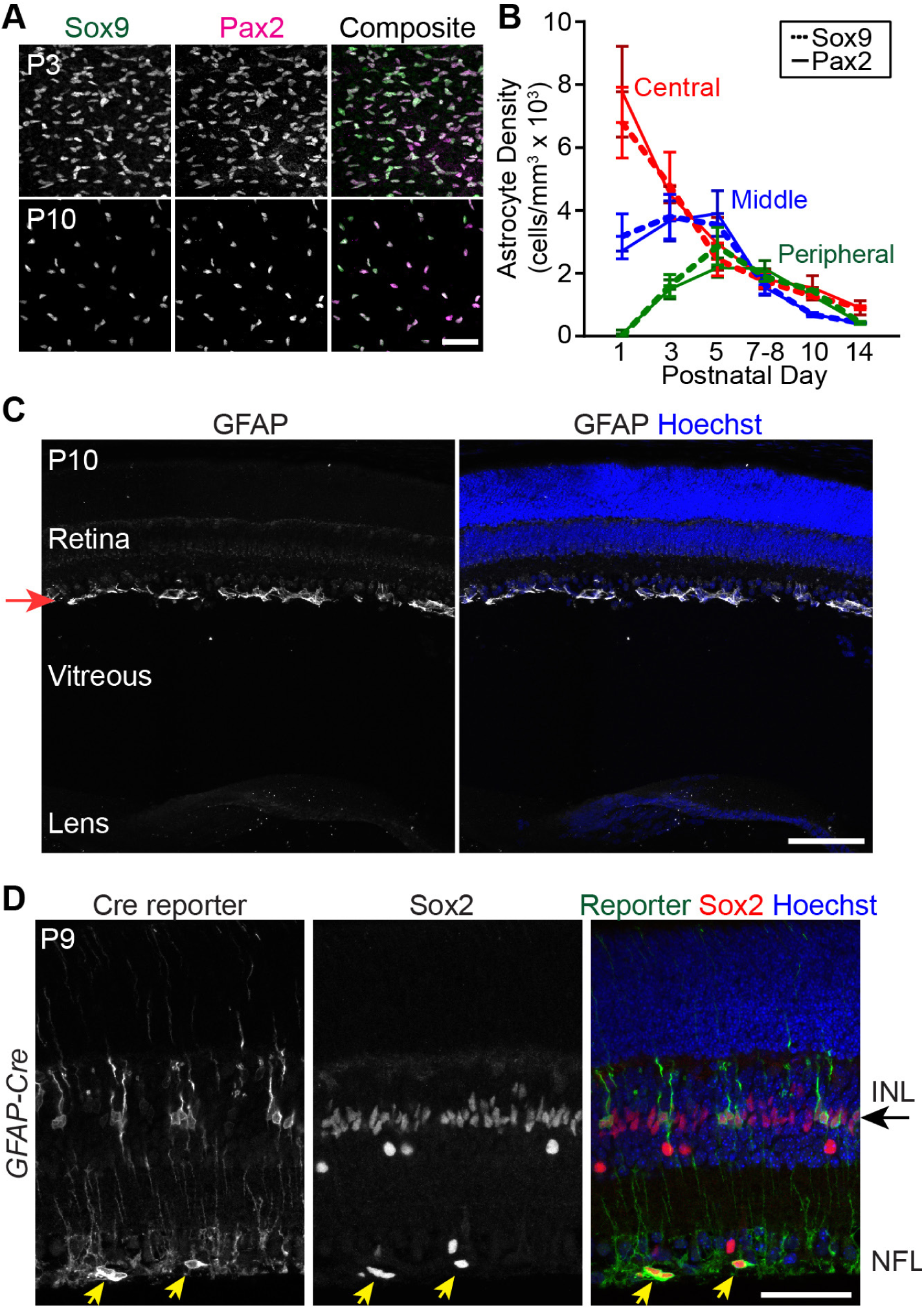
Evaluation of possible mechanisms underlying developmental astrocyte loss. **A,B**) Astrocyte loss confirmed using two independent markers, Sox9 and Pax2. A: Confocal images of RNFL imaged *en face* in whole-mount retinas. Pax2 and Sox9 colocalize in virtually all astrocyes, at each of two timepoints, demonstrating that both markers label the complete astrocyte population across development. B: Quantification of Sox9^+^ and Pax2^+^ astrocyte densities in central, middle and peripheral retina across development. Density dynamics for each form of astrocyte labeling are nearly identical, suggesting that astrocyte loss cannot be explained by developmental downregulation of Pax2. Sample sizes: Pax2: N = 3 (P1, 5, 7-8, 10, 14) or N = 4 (P3); Sox9: N = 2 (P1, 5), N = 3 (P10, 14), N = 4 (P7-8), or N = 5 (P3). **C**) Astrocytes do not migrate out of the RNFL during the death period. Whole-eye cross-section from P10 mouse, stained for GFAP to label astrocytes, and counterstained with Hoechst. Astrocytes are only found within the retina (in the RNFL; red arrow), and have not migrated into extra-retinal spaces such as the vitreous or lens. P10 was chosen for this analysis because astrocyte numbers have declined substantially by this age, so if migration was a major cause of astrocyte loss we should have seen many astrocytes in non-retinal regions by this time. **D**) Astrocyte lineage tracing during the period of astrocyte loss, using *GFAP-cre*mice crossed to a Cre reporter (*Rosa26*^*iDTR*^). P9 retinal cross-sections were immunostained with anti-DTR to reveal cells that experienced Cre activity, and for Sox2 as a marker of astrocytes and Müller glia. Cre reporter expression is only found within two astrocytic cell types: Sox2^+^ astrocytes of the RNFL (yellow arrows), and Sox2^+^ Müller glia within the INL (black arrow). This finding demonstrates that GFAP^+^ astrocytes do not transdifferentiate into a non-astrocytic cell type. Since astrocyte precursors activate Cre expression as early as P0-1 [30], we would expect to see reporter-positive neurons by P9 if transdifferentiation were responsible for the decline in astrocyte numbers. If RNFL astrocytes were trandsifferentiating into Müller glia, we would expect to see some reporter-positive cells migrating between the RNFL and the INL, where Müller cells reside (black arrow). However, no migrating reporter-positive cells were observed. Error bars: mean ± S.D. Scale bars: 50 µm (A,D); 100 µm (C).

**Figure S2.**
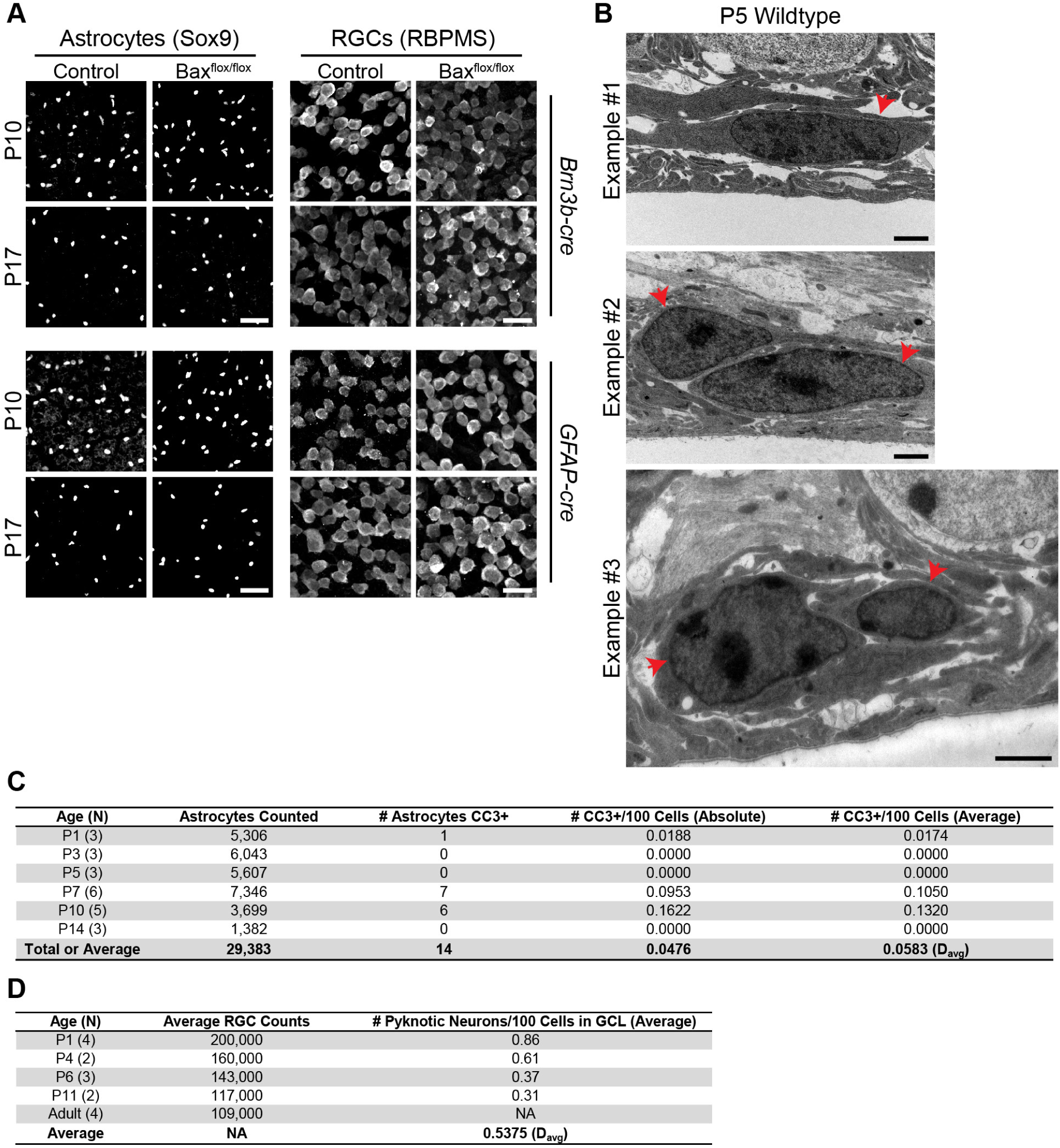
Assessment of astrocyte apoptosis. **A**) Confocal images illustrating astrocyte and RGC densities in control and *Bax* mutant mice. Images similar to these were used for quantification shown in Fig. 2D,E. Sox9^+^ astrocytes did not differ in density between wildtype controls and cell-type-specific *Bax* mutants (left panels). More RBPMS^+^ RGCs are evident following *Bax*deletion in RGCs (*Brn3b-cre*), but not following *Bax*deletion in astrocytes (*GFAP-Cre*). **B**) Representative electron micrographs showing ultrastructural morphology of retinal astrocytes at P5. Red arrows = astrocyte nuclei. Note absence of anatomical features typical of various cell death pathways, such as: 1) apoptosis (condensed nuclei); 2) necrosis (swollen cells/nuclei); 3) autophagy (vacuolization)[33]. Analysis was performed on 2 sections from each of 5 animals; overall, 69 images were analyzed, each of which typically contained 1-3 astrocytes, and occasionally contained >3 astrocytes. **C**) Quantification of retinal astrocytes expressing CC3 across development. ***Davg*** = average death rate (utilized in model found in Fig 2C; see Methods). Overall values for the columns “Astrocytes Counted” and “# Astrocytes CC3+” are totals; overall values for the columns “#CC3/100 Cells (Absolute)” and “#CC3/100 Cells (Average)” are averages. **D**) Data from Perry et al. (1983) quantifying rat RGCs and the number of pyknotic GCL neurons across development. These data were utilized in model found in Fig 2B. ***Davg*** = average death rate (see Methods). Scale bars: 50 µm (A, Astrocytes); 25 µm (A, RGCs); 2 µm (B).

**Figure S3.**
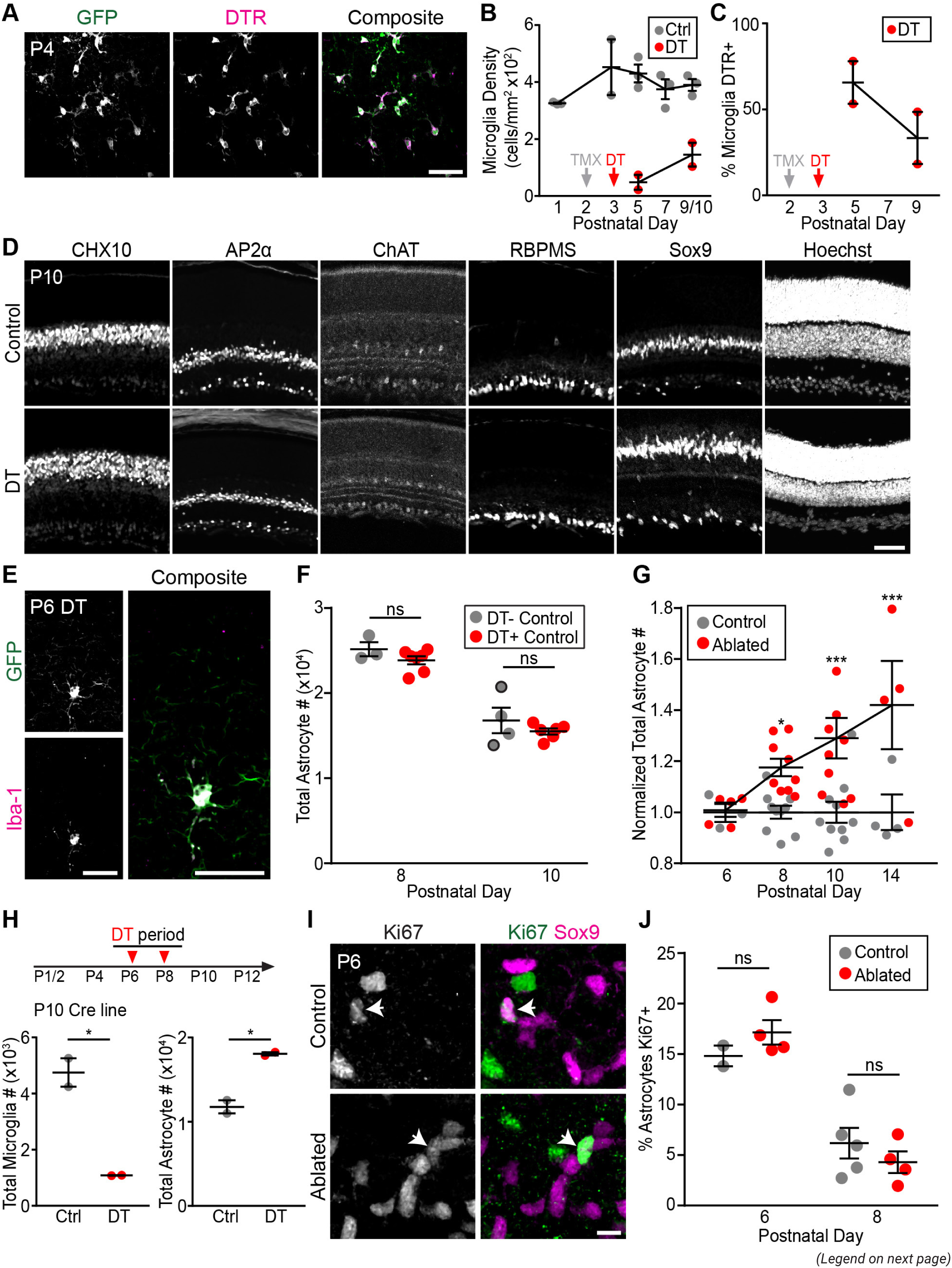
Ablation of microglia using inducible DTR system increases astrocyte number. **A)** Representative image of microglia from P4 *Cx3cr1*^*CreER-ires-YFP*^*;Rosa26*^*iDTR*^ retina, stained for anti-GFP and anti-DTR. Mice received one dose of TMX at P2 to induce expression of DTR. Virtually all GFP^+^ microglia are also DTR^+^. See Results for cell count data. **B)** Quantification of RNFL microglia density following a single round of TMX and diphtheria toxin (DT), administered at the indicated timepoints (gray, red arrows). In *Cx3cr1*^*CreER*^*; Rosa26*^*iDTR*^ animals (red data points), microglia were largely eliminated by 2 days post DT, but significant repopulation was seen by 4-5 days post-DT. Based on this finding, we administered DT at two-day intervals in our long-term ablation paradigm (Fig. 6B). Gray data points: Control data from non-littermate animals from the *Cx3cr1*^*CreER-ires-YFP*^ background for comparison; these animals did not receive TMX or DT. **C)** Quantification of DTR expression by spared microglia in the same ablated animals shown in B. At 2 days post-DT, few microglia remain (B), but a substantial fraction of these are DTR-negative. The DTR-negative fraction is even higher by 6 days post-injection, suggesting that much of the repopulation is performed by microglia that escaped CreER-mediated DTR expression. This finding led us to conclude that long-term microglia ablation would require multiple TMX injections (as in the paradigm described in Fig. 6B). **D)** Representative retinal cross-sections from P10 DT ablated mice or their littermate controls. Microglial ablation was performed following the paradigm described in Fig. 6B. Staining for the major retinal cell types shows that overall retinal histology appears largely normal in ablated retinas. Antibodies used: CHX10 for bipolar cells; AP2α for amacrine cells; ChAT for starburst amacrine cells (also shows sublaminar integrity of IPL); RBPMS for RGCs; Sox9 for *Müllergliaandastrocytes.Hoechstservedasnuclearcounterstain.* **E)** Two microglia-selective markers, *Cx3cr1*^*CreER-ires-YFP*^and Iba-1, confirm absence of retinal microglia in DT ablated animals. Field of view was chosen to show a single microglial cell that escaped DT-mediated ablation. This cell is co-stained by antibodies to GFP and Iba1; however, no other cells in the field of view are positive for either marker. Ablation was via the TMX/DT paradigm described in Fig. 6B. Note that a third marker, CD45, also confirmed absence of microglia (see Fig. 7E; S5 Fig. 5E). **F)** No difference in astrocyte number between control mice that received DT injections (red) and those that did not receive DT (gray). In early experiments, our breeding strategy was such that all mice in the litter inherited both CreER and DTR transgenes. In these cases, the control mice received TMX but not DT (except in 2 cases, where control animals received neither TMX or DT – data for these animals is denoted by gray dots with black outlines). For subsequent experiments we changed our breeding strategy so that some mice would inherit only one of the two transgenes, allowing us to administer TMX and DT to all animals and still obtain unablated controls. To ask whether these two types of controls were equivalent (DTvs DT+), astrocyte numbers were compared at P8 and P10. No significant difference was found (P-values: P8 = 0.5072; P10 = 0.4827), so we pooled both types of controls for subsequent analysis. Statistics: 2-way ANOVA followed by Fisher’s LSD post-hoc test. **G)** Quantification of total astrocyte numbers in control and microglia-ablated retinas. Same data as in Fig. 6D; here the data is plotted normalized to control values, to highlight the magnitude of astrocyte number excess at each age. Statistics: 2-way ANOVA followed by Fisher’s LSD post-hoc test. P-values: P6=0.9508; P8=0.0261; P10=0.0003; P14=0.0008. **H)** Ablation of microglia using BAC transgenic *Cx3cr1-cre* mice (“Cre line”) to drive DTR expression. Microglia were ablated following administration of diphtheria toxin (DT, red arrows) at indicated times (top). Ablated mice carried both *Cx3cr1-cre*and *DTR* transgenes; littermate controls lacked one of the transgenes.Statistics: Two-tailed t-tests. P-values: Microglia, p=0.0185; Astrocytes, p=0.0158. **I,J**) Ki67 immunostaining was used to assess astrocyte proliferation following microglia ablation (TMX/DT paradigm described in Fig. 6B). I: Representative images of Ki67^+^ cells in RNFL at P6. Examples of Ki67^+^/Sox9^+^ astrocytes are shown (arrows). J: Quantification of astrocyte proliferation at P6 and P8. Microglia ablation does not affect the percentage of Ki67^+^ proliferating astrocytes. Statistics: 2-way ANOVA followed by Fisher’s LSD post-hoc test. P-values: P6 = 0.3380; P8 = 0.3202. Error bars: mean ± S.E.M. Sample sizes denoted by data points on graphs. Scale bars: 50 µm (A, D, E); 10 µm (I).

**Figure S4.**
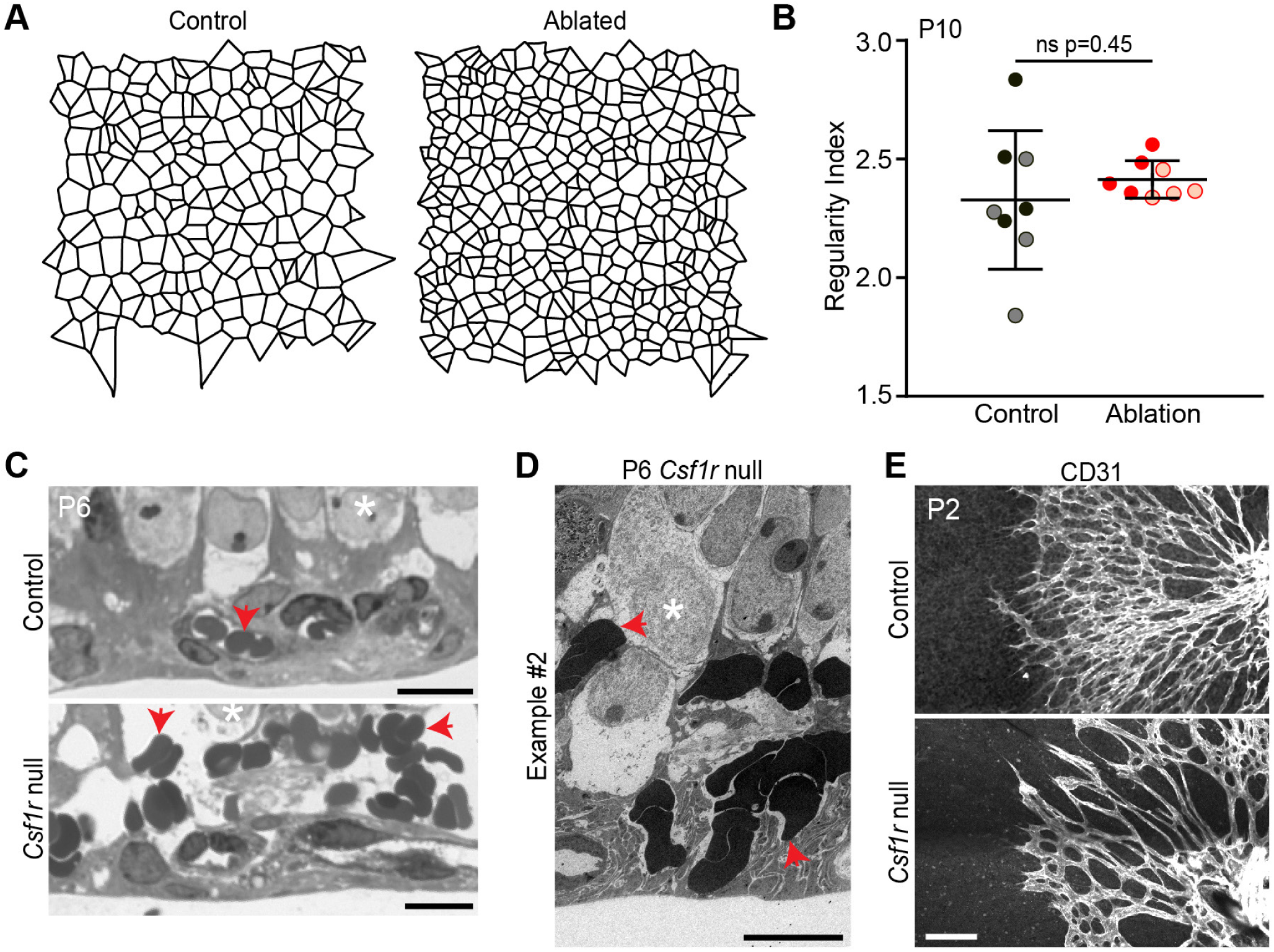
Functional impact of developmental absence of microglia. **A,B**) Excess astrocytes incorporate into the astrocyte mosaic, as shown by Voronoi domain regularity analysis. A: Representative Voronoi domains for control and DT ablated retinas at P10. Voronoi domains show the set of points nearest to each astrocyte within the cellular array. Cell types that show local cell-cell repulsion, such as astrocytes, will have Voronoi domains of fairly uniform size, as is the case for the control astrocyte array (left). In ablated animals, Voronoi domains are smaller due to the increase number of astrocytes. However, the size of the domains are still fairly uniform, suggesting that the increased cell number has not affected array regularity. B: Voronoi domain regularity indices (i.e mean / S.D. of the domain areas) were calculated for the astrocyte arrays in control and ablated animals. Points, individual measurements; shading denotes different animals. There is no change in regularity between the two groups. If excess astrocytes in ablated animals were dead, we would expect they would be randomly distributed and thus would lower the regularity index. As this was not observed we conclude the excess astrocytes are incorporated normally into the non-random mosaic pattern. **C**) Higher magnification view of the same control and *Csf1r* mutant thin plastic sections shown in Fig. 6I, to more clearly show that control RBCs are inside vasculature. Red arrows = red blood cells. White asterisks = RGCs. **D**) An additional example of a representative electron micrograph from *Csf1r* mutant retina. Red arrows indicate examples of extravascular red blood cells which have accumulated in NFL-GCL region, indicative of bleeding. White asterisks = RGCs. **E**) Overall anatomy of CD31^+^ retinal vasculature at P2 in control and *Csf1r* null retinas. Consistent with previous reports [58], animals lacking microglia have somewhat reduced branching frequency at P2. These are not maintained at later ages ([58] and data not shown). Error bars: mean ± S.D. Scale bars: 10 µm (C, D); 100 µm (E).

**Figure S5.**
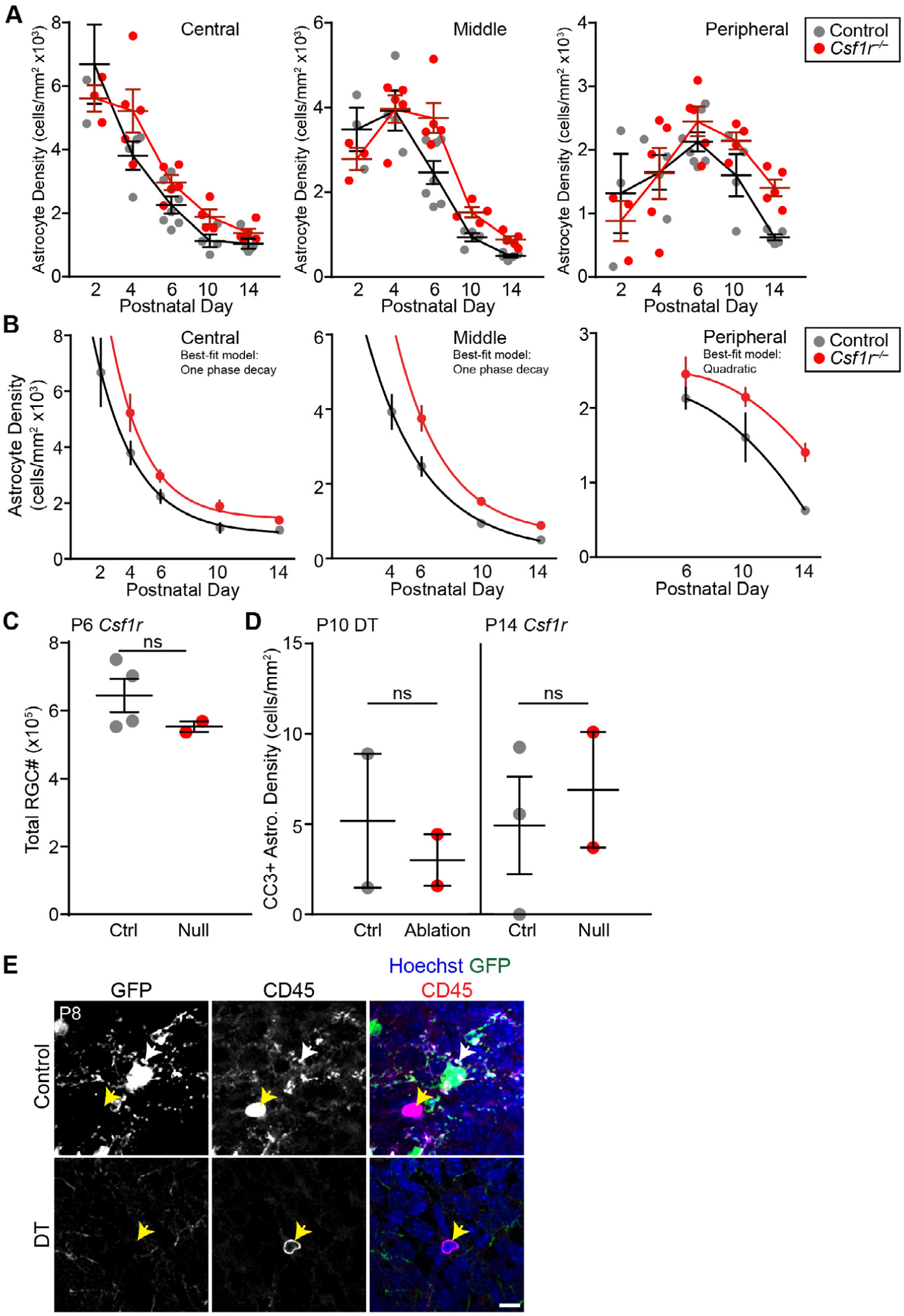
Assessment of compensatory mechanisms driving astrocyte death. **A)** Temporal dynamics of astrocyte density in *Csf1r* mutants (red) and littermate controls (black). In controls, a rising phase (due to cell addition; see Fig. 1A) is followed immediately by a decline phase due to microglia-mediated cell death. Mutant curves are right-shifted due to delayed onset of decline phase – i.e. delayed onset of compensatory non-microglial death. Decline phase dynamics are similar between genotypes (also see B). Curves were compared by nonlinear regression analysis (see B for details). **B**) Nonlinear regression analysis of the *Csf1r* mutant and littermate control astrocyte density curves shown in A. The decline phase of each curve from each regional dataset was modeled (i.e. Central: P2-P14 (control), P4-P14 (null); Middle: P4-P14 (control), P6-P14 (null); Peripheral: P6-P14 (control and null)). A one-phase exponential decay model best-fit Central and Middle curves, while a second order polynomial (quadratic) model best-fit Peripheral curves. Model parameters were similar for each mutant-control pair (Central: t=2.789 control, t=2.331 mutant; Middle: t=3.947 control, t=3.237 mutant; Peripheral: B0=2066, B1=95.04, B2=-14.13 control; B0=2089, B1=141.4, B2=-13.59 mutant). **C**) Quantification of total RGC numbers at P6 in control and *Csf1r* null retinas. RGC numbers are unchanged in the absence of microglia. This suggests that RGC death proceeds normally in *Csf1r* mutants and that loss of microglia cannot rescue them from death. Further, it suggests that increased CC3^+^ nuclei (Fig. 7D) are a result of corpse accumulation rather than excessive RGC death. Statistics: Two-tailed t-test (p = 0.2835). **D**) CC3^+^ astrocyte density at P10 or P14 in DT ablated or *Csf1r* mutant retinas. Each ablation paradigm was compared to its respective littermate controls. Density of CC3^+^ astrocytes was not increased in either paradigm at these timepoints, further supporting the conclusion from Fig. 7B that absence of microglia does not increase astrocyte apoptosis rates. Statistics: Two-tailed t-test (p-values: DT=0.6396; *Csf1r* = 0.6728). **E**) Representative images of CD45^+^ cells at P8 in control and DT ablated retinas. Images similar to these were used for quantification shown in Fig. 7E. White arrow: GFP^+^/CD45^+^ microglia. Yellow arrow: GFP^−^/CD45^+^ leukocyte. Error bars: mean ± S.E.M. Sample sizes denoted by data points on graphs (note that in (A) “Central” graph is missing one control data point located outside of y-range). Scale bar: 10 µm.

### Supplementary Movies

#### Movie S1. Astrocyte debris within microglia

Three-dimensional reconstruction of confocal Z stack depicted in Fig. 4A. Microglia (green) contain Pax2+ astrocyte debris (red; yellow in overlay). Reconstruction shows that debris is genuinely located inside the microglial cells. Microglia labeled by anti-GFP in *Cx3cr1*^*CreER*^ animals.

#### Movie S2. Partial enclosure of astrocytes by microglial cells

Three-dimensional reconstruction of a confocal Z stack showing microglia (anti-GFP, green) interacting with astrocytes (Pax2, red). At center of image, an astrocyte nucleus is partially enclosed by numerous arbors from a microglial cell. At bottom of image (at right in first frame), a microglial cell partially encloses two astrocytes; this cell also contains Pax2^+^ debris (yellow).

#### Movie S3

Extensive contact between microglial cells and astrocytes. Three-dimensional reconstruction of a confocal Z stack showing three microglial cells (anti-GFP, green) interacting extensively with multiple astrocyes. For the purposes of Fig. 4D, these interactions were largely scored as “touching.”

