## Supplementary figures and images for "Large-scale death of retinal astrocytes during normal development mediated by microglia"

### S1 Movie

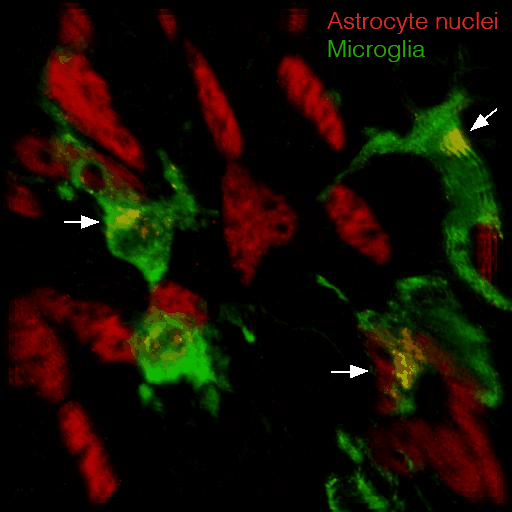

### S3 Movie

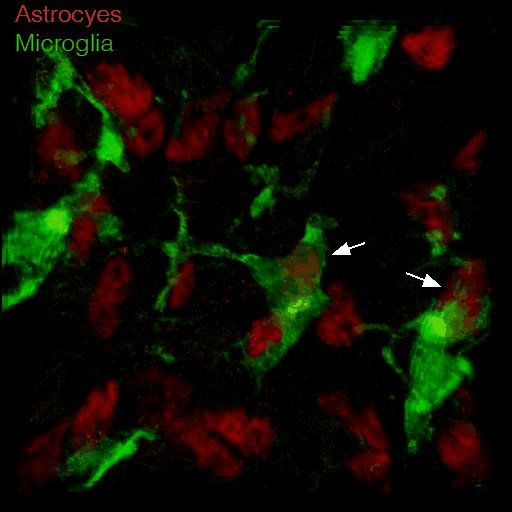
